# Glycogen fuels and spatially regulates nucleotide biosynthesis in cancer

**DOI:** 10.64898/2026.11.21.768289

**Authors:** Tashbib Khan, Stefan A. Harry, Mario C. Chang, Thasin Peyear, Grace Figlioli, Marjan Berishaj, Madeline Gish, Reece Larson, Jonah Lee, John M. Asara, John G. Clohessy, Hadrien Dyvorne, Myriam M. Chaumeil, Matthew S. Gentry, Kayvan R. Keshari, Liron Bar-Peled, Ramon C. Sun, Alex Toker

## Abstract

Cancer cells must coordinate metabolic substrate availability with the organization of biosynthetic pathways to sustain proliferation. Glycogen accumulates in many cancers, but its function extends beyond glucose storage. Here, we show that glycogen regulates nucleotide biosynthesis through distinct metabolic and spatial mechanisms. Disrupting glycogenolysis impairs tumor growth and mitochondrial pyruvate oxidation, compromises NADPH homeostasis, and selectively reduces glutamine utilization for reductive biosynthesis. Cytoplasmic and lysosomal glycogenolysis produce different defects in nucleotide precursor metabolism and distinct dependencies on biosynthetic enzymes. By contrast, disrupting glycogen synthesis creates a selective vulnerability to depletion of CAD, the multifunctional enzyme rate limiting for pyrimidine biosynthesis. Pyrimidine intermediates associate with glycogen particles, while glycogen availability influences their spatial distribution and the organization of the pyrimidine biosynthetic enzyme UMP synthase. Together, these findings identify glycogen as a source of biosynthetic substrates and a regulator of nucleotide metabolic organization, linking carbon storage and mobilization to cancer cell proliferation.

**HIGHLIGHTS:**

- Glycogen breakdown is required for tumor growth and mitochondrial oxidation in vivo
- Glycogenolysis fuels reductive glutamine metabolism, not energy, in cancer cells
- Cytoplasmic and lysosomal glycogen breakdown fuel distinct nucleotide pathways
- Perturbing glycogen metabolism alters pyrimidine chanelling

**“IN BRIEF”:** Khan et al. identify a dual role for glycogen in cancer cell nucleotide biosynthesis. Cytoplasmic and lysosomal glycogenolysis support mitochondrial biosynthetic metabolism and provide distinct inputs into nucleotide synthesis. The glycogen particle itself influences pyrimidine metabolism by regulating the spatial organization of biosynthetic intermediates and enzymes.

## INTRODUCTION

Glycogen is a highly branched glucose polymer that serves as a metabolic reserve, supplying glucose to meet acute energetic demands in tissues such as skeletal muscle and liver^1,2^. Although glycogen accumulation is a common feature of cancer^3,4^, its contribution to tumor metabolism remains poorly understood. Glycogen synthesis and degradation are regulated by oncogenic signaling pathways, including p53^5^ and PI3K/AKT^6,7^, and by metabolic stresses such as hypoxia^8,9^. These observations suggest that glycogen metabolism may serve functions beyond maintaining cellular energy homeostasis, particularly in cancer cells that must coordinate metabolic substrate availability with the biosynthetic demands of proliferation^10^. However, whether glycogen primarily provides carbon for energy production or supports other metabolic processes remains unclear.

The contribution of glycogen to cancer progression has been challenging to define. Studies in clear cell renal cancer have reported conflicting effects of glycogen metabolism on tumor growth and therapeutic resistance^9,11^, suggesting that its function may depend on cellular or metabolic context. Glycogen abundance is also highly heterogeneous within tumors, and the pathways governing its turnover are functionally distinct. Cytoplasmic glycogenolysis, mediated by glycogen phosphorylase and glycogen debranching enzyme, mobilizes glycogen independently of its lysosomal degradation by acid α-glucosidase. Whether these pathways make equivalent contributions to cancer cell metabolism, or support distinct metabolic processes, is unknown. *De novo* nucleotide biosynthesis requires the coordinated supply of ribose phosphate, glutamine-derived nitrogen, aspartate, and one-carbon units from interconnected cytoplasmic and mitochondrial metabolic pathways^12,13^. Whether glycogen metabolism contributes to the availability of these biosynthetic precursors remains unclear. Moreover, glycogen may have functions independent of its degradation. Recent studies have demonstrated that glycogen particles can regulate signaling through the sequestration of Hippo pathway components, thereby promoting YAP-dependent liver tumorigenesis^14^. These findings raise the possibility that glycogen influences cellular metabolism not only through the release of stored glucose, but also through the spatial organization of metabolic pathways.

Here, we identify complementary metabolic and spatial functions for glycogen in cancer cell nucleotide biosynthesis. Using genetic perturbations, isotope tracing, in vivo hyperpolarized ^13^C spectroscopy, and spatial metabolic analyses, we demonstrate that glycogenolysis supports mitochondrial pyruvate oxidation, NADPH homeostasis, and reductive glutamine metabolism. We further show that cytoplasmic and lysosomal glycogenolysis make distinct contributions to nucleotide precursor metabolism and establish different dependencies on nucleotide biosynthetic enzymes. Beyond these metabolic functions, we find that the glycogen particle associates with pyrimidine biosynthetic intermediates and influences the spatial organization of UMP synthase. These findings establish glycogen as both a source of biosynthetic substrates and a regulator of nucleotide metabolic organization, linking glycogen storage and turnover to the metabolic requirements of cancer cell proliferation.

## RESULTS

### Cytoplasmic glycogenolysis supports mitochondrial pyruvate oxidation in tumors

Glycogen accumulates in many human cancers, although its distribution within tumors is highly heterogeneous^1,3,15–17^. To investigate the spatial relationship between glycogen metabolism and other metabolic pathways, we analyzed publicly available spatial transcriptomic datasets from breast, lung, and colon cancers. We generated a glycogenolysis signature comprising genes encoding the cytoplasmic glycogen phosphorylase enzymes (*PYGL* and *PYGB*), glycogen debranching enzyme (*AGL*), and lysosomal acid α-glucosidase (*GAA*), and identified the 100 genes most strongly correlated with this signature across tumor sections (**Figure 1A**). Pathway enrichment analysis identified glycolysis, as expected, but also revealed associations with mitochondrial carbon metabolism, including pyruvate entry into the tricarboxylic acid (TCA) cycle, and pyrimidine nucleotide metabolism (**Figure 1B**). These findings suggested that glycogenolysis may support metabolic processes extending beyond its established role in supplying glucose for glycolysis.

**Figure 1.**
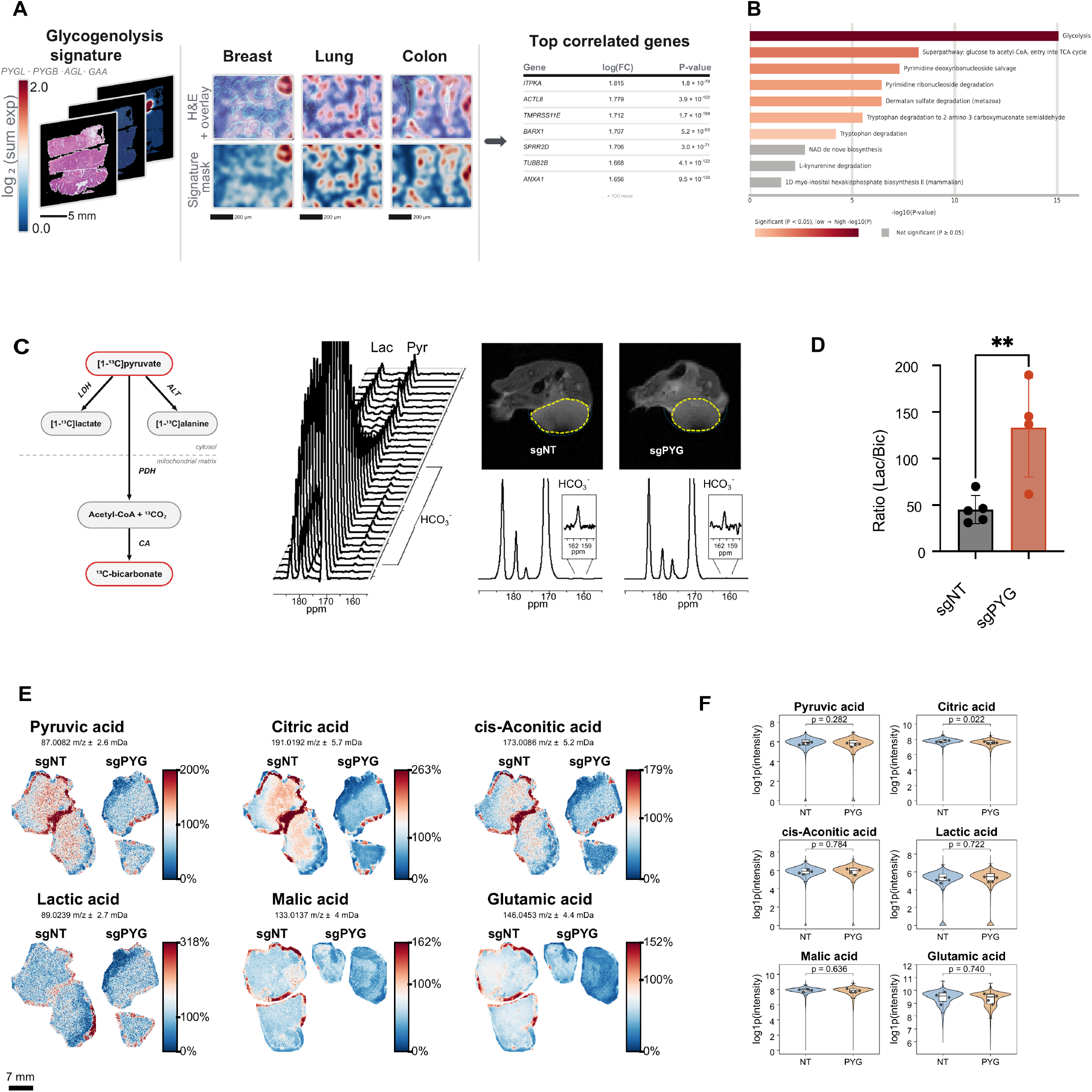
Cytoplasmic glycogenolysis supports mitochondrial pyruvate oxidation and tumor growth in vivo. **(A)** Spatial transcriptomics workflow. Visium HD sections of breast, lung, and colon cancer were scored for a glycogenolysis signature (*PYGL*, *PYGB*, *AGL*, and *GAA*; log_2_ summed expression), shown as H&E overlays and signature masks. The 100 genes most correlated with the signature across sections were identified (right; top-ranked genes with log fold change and p value). **(B)** Pathway enrichment (Enrichr, HumanCyc 2016) of the 100 signature-correlated genes in (A), ranked by −log_10_(p). Colored bars, p < 0.05; gray bars, not significant. **(C)** Hyperpolarized [1-^13^C]pyruvate magnetic resonance spectroscopy of subcutaneous sgNT and sgPYG SUM159PT xenografts. Left, [1-^13^C]pyruvate is converted to [1-^13^C]lactate and [1-^13^C]alanine by lactate dehydrogenase (LDH) and alanine aminotransferase (ALT) in the cytosol, or to ^13^C-bicarbonate by pyruvate dehydrogenase (PDH) and carbonic anhydrase (CA) in mitochondria. Middle, representative dynamic ^13^C spectra showing pyruvate (Pyr), lactate (Lac), and bicarbonate (HCO_3_^−^) over time. Right, representative T_2_-weighted images (tumor outlined in yellow and surface coil position in dark teal) and summed ^13^C spectra; insets show the expanded bicarbonate region. **(D)** Lactate-to-bicarbonate ratio per tumor (n = 5 sgNT and 4 sgPYG mice). **(E)** Representative MALDI mass spectrometry imaging (MSI) of pyruvic acid (*m/z* 87.0082), citric acid (*m/z* 191.0192), cis-aconitic acid (*m/z* 173.0086), lactic acid (*m/z* 89.0239), malic acid (*m/z* 133.0137), and glutamic acid (*m/z* 146.0453) in orthotopic sgNT and sgPYG SUM159PT xenografts. Intensities are TIC normalized and scaled to the mean. Scale bar, 7 mm. **(F)** Pixel-level log1p(TIC-normalized intensity) for the metabolites in (E) (n = 4 tumor sections per genotype). Violins show all pixels; boxes show median and interquartile range (whiskers, 1.5× IQR); points show per-section means. p values were calculated with a linear mixed-effects model with tissue section as a random intercept. Data in (D) are represented as mean ± SD; two-tailed unpaired t test, \*\**p* < 0.01. sgNT, non-targeting control; sgPYG, *PYGB*/*PYGL* double knockout. See also Figure S1.

To determine whether glycogenolysis contributes to mitochondrial metabolism *in vivo*, we disrupted cytoplasmic glycogen breakdown by deleting both PYGB and PYGL, which encode functionally redundant glycogen phosphorylase isoforms^9,18^. CRISPR-Cas9-mediated deletion of *PYGB* and *PYGL* (sgPYG) in SUM159PT breast cancer cells significantly impaired orthotopic tumor growth in NSG mice, as determined by longitudinal tumor growth measurements and endpoint tumor weights, without appreciable changes in tumor histology (**Figures S1A-S1E**). To assess mitochondrial metabolism directly, we performed hyperpolarized [1-^13^C]pyruvate magnetic resonance spectroscopy on awake mice before substantial differences in tumor size had emerged. This approach permits the noninvasive measurement of pyruvate metabolism *in vivo*, including its conversion to lactate by lactate dehydrogenase and to bicarbonate following mitochondrial pyruvate oxidation by pyruvate dehydrogenase^19–21^ (**Figure 1C**). Scanning animals under awake conditions overcomes the metabolically and physiologically suppressive effects of anesthesia and maximizes the production of lactate and bicarbonate from pyruvate which is essential for detecting decrements from baseline levels^22^. Although pyruvate-derived lactate and alanine signals were not significantly altered, bicarbonate production was markedly reduced in sgPYG tumors (**Figure S1F**). Accordingly, the lactate-to-bicarbonate ratio increased nearly threefold following loss of cytoplasmic glycogenolysis (**Figure 1D**). These findings indicate that glycogenolysis supports mitochondrial pyruvate oxidation in tumors, independently of the subsequent effects on tumor growth.

We next examined whether impaired pyruvate oxidation was accompanied by alterations in the abundance and spatial distribution of mitochondrial metabolites. Matrix-assisted laser desorption/ionization mass spectrometry imaging (MALDI-MSI) of sgNT and sgPYG tumors revealed reduced signals for several TCA cycle-associated metabolites, including citrate, cis-aconitate, and malate, following disruption of glycogenolysis (**Figures 1E and S1G**). Despite considerable intratumoral heterogeneity, citrate abundance was significantly reduced in sgPYG tumors, whereas differences in pyruvate, lactate, cis-aconitate, malate, and glutamate did not reach statistical significance across tumors (**Figure 1F**). Together with the hyperpolarized pyruvate measurements, these findings establish a functional relationship between cytoplasmic glycogenolysis and mitochondrial carbon metabolism *in vivo*.

### Cytoplasmic and lysosomal glycogenolysis sustain mitochondrial carbon metabolism under nutrient stress

Glycogen is degraded through two distinct pathways: cytoplasmic glycogenolysis, mediated by glycogen phosphorylase and glycogen debranching enzyme, and lysosomal glycogenolysis, mediated by glycophagy and acid α-glucosidase (GAA) (**Figure 2A**). To determine the contribution of each pathway to mitochondrial metabolism, we disrupted cytoplasmic glycogenolysis by deleting *PYGB* and *PYGL* (sgPYG), or lysosomal glycogenolysis by deleting *GAA* (sgGAA), in SUM159PT breast cancer, SKOV3 ovarian cancer, PC3 prostate cancer, and 143B osteosarcoma cells (**Figures S2A and S2C**). Anti-glycogen immunofluorescence^23^ revealed substantial glycogen accumulation in SUM159PT and SKOV3 cells following disruption of either pathway (**Figure 2B**), consistent with the contribution of both cytoplasmic and lysosomal degradation to glycogen turnover. Although glycophagy has traditionally been associated with the clearance of abnormal glycogen, recent studies have implicated this pathway in hepatic glucose production and lung cancer growth^24,25^.

**Figure 2.**
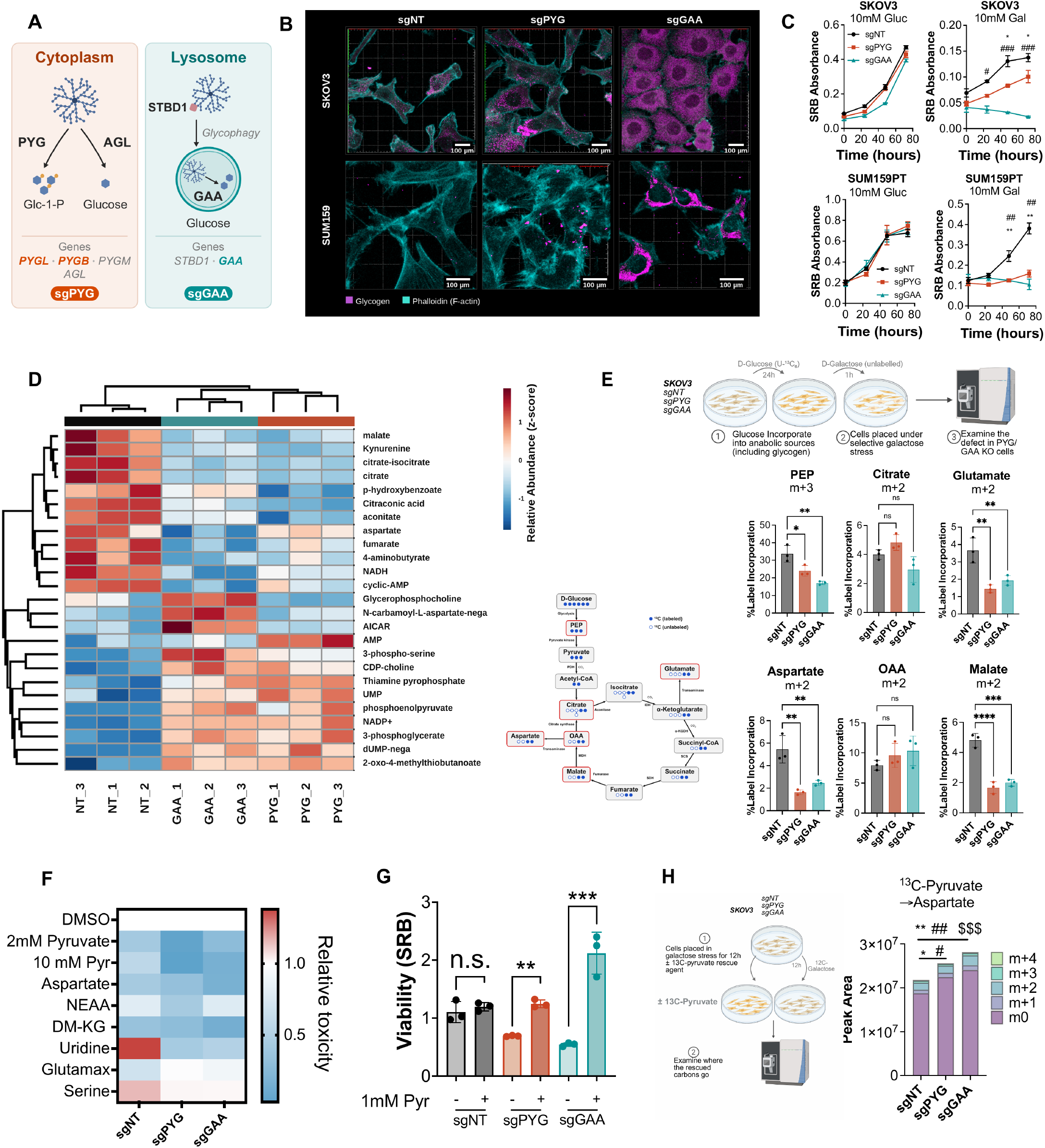
Cytoplasmic and lysosomal glycogenolysis support mitochondrial carbon metabolism and aspartate production under nutrient stress. **(A)** Cytoplasmic glycogenolysis is mediated by glycogen phosphorylase (PYGB, PYGL, and PYGM) and glycogen debranching enzyme (AGL); lysosomal glycogenolysis is mediated by STBD1-dependent glycophagy and acid α-glucosidase (GAA). **(B)** Representative immunofluorescence of glycogen (magenta) and F-actin (phalloidin, cyan) in SKOV3 (top) and SUM159PT (bottom) cells. Scale bars, 100 μm. **(C)** Growth of SKOV3 (top) and SUM159PT (bottom) cells in 10 mM glucose (left) or 10 mM galactose (right), measured by sulforhodamine B (SRB) absorbance (n = 3). * and # denote sgNT versus sgPYG and sgNT versus sgGAA, respectively. **(D)** Top 25 altered polar metabolites in SKOV3 cells after 1 h in galactose (log fold change, row normalized, hierarchical clustering). **(E)** Glycogen pulse-chase. SKOV3 cells were labeled with U-^13^C-glucose for 24 h and chased into unlabeled galactose for 1 h (top). Label incorporation into the circled TCA cycle intermediates (left) is shown at right: PEP m+3 and citrate, glutamate, aspartate, oxaloacetate (OAA), and malate m+2. **(F)** Relative galactose toxicity, normalized to DMSO, with the indicated supplements: pyruvate (2 or 10 mM), aspartate, non-essential amino acids (NEAA), dimethyl-α-ketoglutarate (DM-KG), uridine, GlutaMAX, or serine. **(G)** SRB absorbance of SKOV3 cells in galactose with or without 1 mM pyruvate. **(H)** Aspartate isotopologues (m+0 to m+4) in cells cultured in galactose with U-^13^C-pyruvate (schematic, left). *, #, and $ denote comparisons of m+0, m+1, and m+2, respectively. Data are represented as mean ± SEM; n = 3 biological replicates unless otherwise stated; one-way ANOVA with multiple comparisons (\**p* < 0.05, \*\**p* < 0.01, \*\*\**p* < 0.001, \*\*\*\**p* < 0.0001; ns, not significant). sgNT, non-targeting control; sgPYG, *PYGB*/*PYGL* double knockout; sgGAA, *GAA* knockout. See also Figures S2, S3, and S4.

Under glucose-replete conditions, disruption of either glycogenolytic pathway had little effect on cell proliferation. By contrast, replacing glucose with galactose, which increases dependence on mitochondrial oxidative metabolism^26–28^, markedly impaired the growth of sgPYG and sgGAA cells across all four cancer cell lines (**Figures 2C and S2B**). A similar dependence on glycogenolysis was observed following glucose withdrawal (**Figure S2D**). Pharmacological inhibition of glycogen phosphorylase with CP91149 ^18,29^ also selectively impaired survival in galactose-containing medium (**Figure S2E**). Moreover, combined inhibition of glycogen phosphorylase and glucose uptake, using CP91149 and the GLUT1 inhibitor BAY-876^30,31^, reduced cell viability across six cancer cell lines (**Figure S2F**). Thus, both cytoplasmic and lysosomal glycogenolysis become important for cancer cell survival when glucose availability or utilization is restricted.

To define the metabolic consequences of impaired glycogenolysis, we performed metabolomic profiling of SKOV3 and SUM159PT cells following acute galactose exposure. Glycogen stores were mobilized within 1 h of galactose treatment (**Figure S3B**), providing a time point to examine the metabolic consequences of glycogen breakdown. Loss of either glycogenolytic pathway reduced the abundance of several TCA cycle intermediates, including citrate, aconitate, fumarate, and malate, accompanied by alterations in glycolytic intermediates and other metabolites (**Figures 2D and S3A**). To determine whether these changes reflected altered utilization of glycogen-derived carbon, we pulse-labeled glycogen with U-^13^C-glucose^9,16^ for 24 h and subsequently chased the label into unlabeled galactose-containing medium (**Figures 2E and S3C**). Disruption of either cytoplasmic or lysosomal glycogenolysis reduced labeling of phosphoenolpyruvate (PEP m+3), glutamate (m+2), malate (m+2), and aspartate (m+2), whereas labeling of citrate (m+2) and oxaloacetate (m+2) was largely preserved (**Figure 2E**). Pharmacological inhibition of glycogen phosphorylase produced a similarly selective alteration in mitochondrial carbon labeling in SKOV3, SUM159PT, and A549 cells (**Figures S3D-S3F**). These findings indicate that glycogenolysis supports specific components of mitochondrial carbon metabolism rather than uniformly supplying carbon to the TCA cycle.

We next determined whether the metabolic defects caused by impaired glycogenolysis could be rescued by supplying alternative carbon sources or biosynthetic precursors. Supplementation with pyruvate, aspartate, or dimethyl-α-ketoglutarate partially rescued cell survival in galactose, whereas uridine, nonessential amino acids, glutamine, and serine were less effective (**Figure 2F**). Pyruvate produced the strongest rescue of cell growth in both sgPYG and sgGAA cells (**Figure 2G**). To determine how pyruvate restored growth, we traced U-^13^C-pyruvate metabolism under these conditions. Pyruvate supplementation increased the abundance of both labeled and unlabeled aspartate in glycogenolysis-deficient cells, accompanied by alterations in the labeling of additional metabolites (**Figures 2H and S4A**). Because mitochondrial aspartate production is essential for proliferating cells^32,33^, these findings suggest that impaired aspartate availability contributes to the growth defect caused by glycogenolysis disruption. Together, our results establish that cytoplasmic and lysosomal glycogenolysis maintain mitochondrial carbon metabolism under nutrient stress, with an important contribution to aspartate production.

### Glycogenolysis sustains NADPH homeostasis and reductive glutamine metabolism

The selective changes in mitochondrial carbon labeling following glycogenolysis disruption suggested that glycogen-derived carbon contributes differently to oxidative and reductive metabolism. To investigate this, we examined the ratio of m+2 malate to m+2 citrate, an indicator of canonical versus noncanonical TCA cycle metabolism^34–36^. This ratio was significantly reduced in sgPYG and sgGAA SKOV3 cells (**Figure 3A**), with a similar trend in sgGAA SUM159PT cells (**Figure S4B**). Pharmacological inhibition of glycogen phosphorylase with CP91149 also reduced the ratio in SUM159PT and A549 cells (**Figure S4C**). Because these metabolic pathways differ in their redox requirements, we examined whether restoring NAD⁺ or NADPH availability could rescue glycogenolysis-deficient cells. Supplementation with α-ketobutyrate, which promotes NAD⁺ regeneration through lactate dehydrogenase^32^, or 2-deoxy-D-glucose, which can support NADPH production through the oxidative pentose phosphate pathway^37,38^, partially restored growth under galactose stress (**Figures 3B and 3C**). However, expression of alternative oxidase (AOX) or the NADH oxidases LbNOX and mitoLbNOX preferentially supported survival of control cells, with substantially less rescue in glycogenolysis-deficient cells (**Figure S4D**). Direct metabolomic measurements revealed a marked increase in the NADP⁺/NADPH ratio following disruption of either glycogenolytic pathway, whereas the NAD⁺/NADH ratio was largely unchanged (**Figures 3D, S5A, and S5B**). Although glycogenolysis has previously been linked to NADPH production through the pentose phosphate pathway^8,39^, glycogen-derived labeling and abundance of pentose phosphate pathway intermediates were not uniformly reduced (**Figure S4E**). Whole-cell proteomics further revealed changes in enzymes involved in NADPH metabolism, most prominently the cytoplasmic and mitochondrial malic enzymes ME1 and ME2 (**Figure 3E**). Together, these findings associate impaired glycogenolysis with altered NADPH homeostasis rather than a generalized defect in NAD⁺ regeneration.

**Figure 3.**
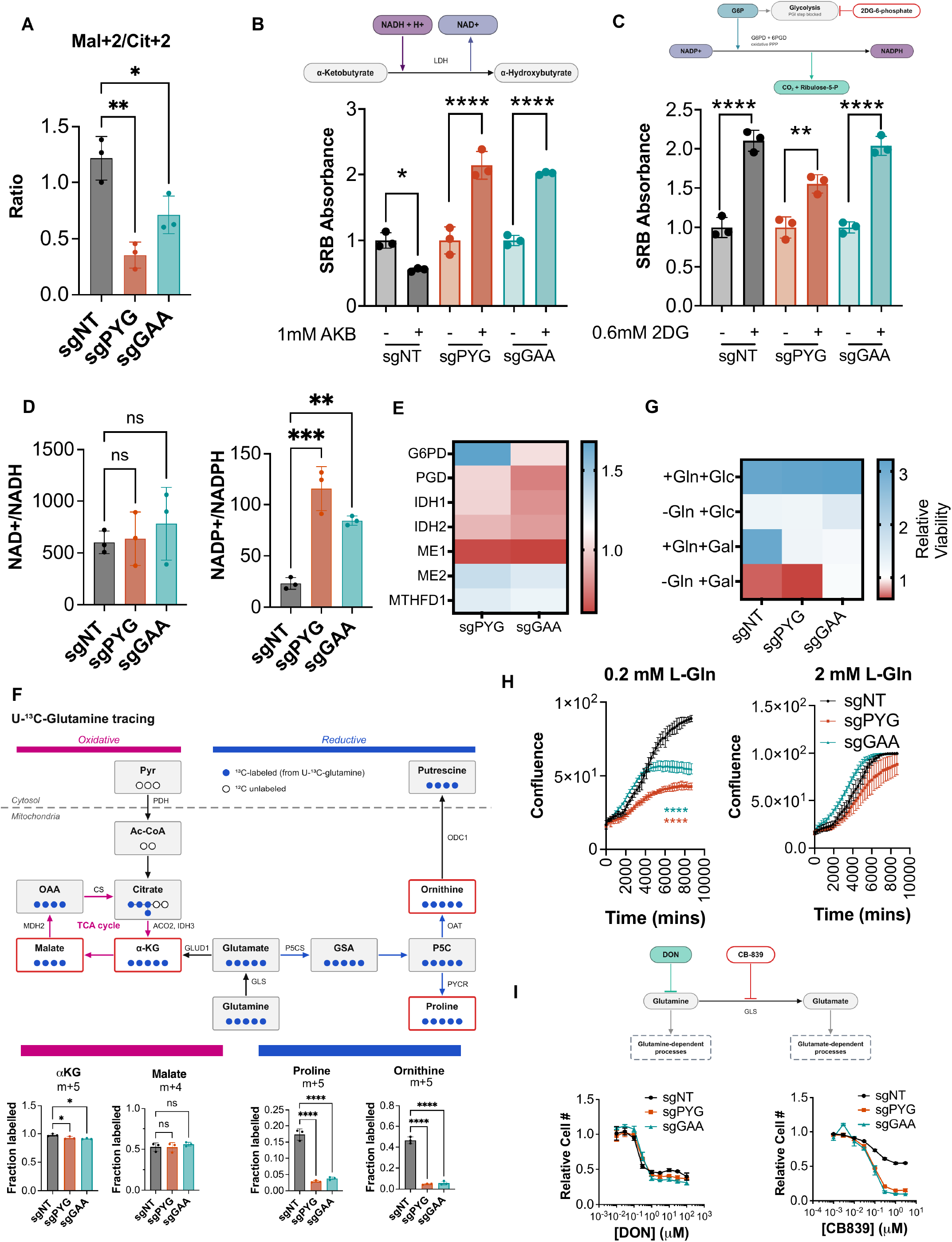
Glycogenolysis maintains NADPH homeostasis and supports reductive glutamine metabolism. **(A)** m+2 malate/m+2 citrate labeling ratio from the tracing in Figure 2E, a readout of canonical versus noncanonical TCA cycle flux. **(B and C)** SRB absorbance of cells in galactose with or without 1 mM α-ketobutyrate (AKB), which regenerates NAD^+^ via lactate dehydrogenase (B), or 0.6 mM 2-deoxy-D-glucose (2DG), which supports NADPH generation via the oxidative pentose phosphate pathway (C). Schematics are shown above. **(D)** NAD^+^/NADH (left) and NADP^+^/NADPH (right) peak-area ratios. **(E)** Whole-cell proteomic abundance of NADPH-generating and one-carbon enzymes in sgPYG and sgGAA cells relative to sgNT (mean of n = 3). **(F)** U-^13^C-glutamine tracing in SKOV3 cells. Oxidative TCA cycle products (α-ketoglutarate [αKG] m+5 and malate m+4) are distinguished from reductive products of P5C metabolism (proline m+5 and ornithine m+5) (top). Fraction labeled is shown below. **(G)** Relative viability of cells cultured with or without glutamine (Gln) in glucose (Glc) or galactose (Gal) (mean of n = 3). **(H)** Confluence over time of cells in 0.2 mM (left) or 2 mM (right) glutamine, measured by live-cell imaging. **(I)** Relative cell number in response to the glutamine antagonist 6-diazo-5-oxo-L-norleucine (DON; left) or the glutaminase (GLS) inhibitor CB-839 (right). Data are represented as mean ± SEM; n = 3 biological replicates; one-way ANOVA with multiple comparisons (\**p* < 0.05, \*\**p* < 0.01, \*\*\**p* < 0.001, \*\*\*\**p* < 0.0001; ns, not significant). sgNT, non-targeting control; sgPYG, *PYGB*/*PYGL* double knockout; sgGAA, *GAA* knockout. See also Figures S4 and S5.

Mitochondrial glutamine metabolism supports both oxidative TCA cycle activity and reductive biosynthetic pathways, including proline and ornithine synthesis^26^. To determine whether glycogenolysis differentially regulates these processes, we traced U-^13^C-glutamine metabolism in SKOV3 and SUM159PT cells. Loss of either glycogenolytic pathway markedly reduced glutamine-derived labeling of proline (m+5) and ornithine (m+5), whereas labeling of malate (m+4) was maintained and α-ketoglutarate (m+5) was only modestly affected (**Figures 3F and S5C**). MALDI-MSI of SUM159PT xenografts similarly revealed reduced proline and ornithine abundance following disruption of cytoplasmic or lysosomal glycogenolysis, with comparatively smaller changes in malate (**Figure S5D**). Consistent with a spatial relationship between glycogen and reductive glutamine metabolism, confocal imaging revealed regions of overlapping glycogen and pyrroline-5-carboxylate synthase (P5CS) signals in SKOV3 cells (**Figure S5E**). These findings indicate that glycogenolysis preferentially supports glutamine utilization for reductive biosynthetic pathways, while largely preserving oxidative glutamine carbon entry into the TCA cycle.

The dependence on glutamine was further examined by manipulating its availability during galactose stress. Glutamine withdrawal largely eliminated the differential survival of control and glycogenolysis-deficient cells in galactose, although viability was markedly reduced across all conditions (**Figure 3G**). Conversely, reducing glutamine availability from 2 mM to 0.2 mM revealed a pronounced growth defect in sgPYG and sgGAA cells that was less apparent under glutamine-replete conditions (**Figure 3H**). Because galactose metabolism increases reliance on glutamine-dependent mitochondrial metabolism^26–28^, these observations suggest that glycogenolysis becomes particularly important when glutamine utilization is limiting.

To distinguish dependence on glutamine itself from its conversion to glutamate, we compared the effects of the glutamine antagonist 6-diazo-5-oxo-L-norleucine (DON)^40^ and the glutaminase inhibitor CB-839^41^. Glycogenolysis-deficient cells displayed similar sensitivity to DON but greater sensitivity to CB-839 than control cells (**Figure 3I**). These findings identify a functional interaction between glycogenolysis and glutaminase-dependent metabolism, consistent with the selective impairment of glutamine-derived reductive biosynthetic products. Collectively, these results establish that cytoplasmic and lysosomal glycogenolysis support NADPH homeostasis and reductive glutamine metabolism, revealing a biosynthetic function for glycogen breakdown that extends beyond its contribution to mitochondrial carbon oxidation.

### Cytoplasmic and lysosomal glycogenolysis differentially regulate nucleotide biosynthesis

The dependence of reductive glutamine metabolism on glycogenolysis prompted us to examine its contribution to nucleotide biosynthesis. Metabolomic profiling of SKOV3 and SUM159PT cells following 6 h of galactose exposure revealed accumulation of nucleotide monophosphates accompanied by reduced abundance of several nucleotide di- and triphosphates in glycogenolysis-deficient cells (**Figures 4A and S6A**). To determine whether these changes reflected altered utilization of glutamine-derived nitrogen, we traced incorporation of amide-^15^N-glutamine into nucleotide precursors. Loss of glycogenolysis increased the unlabeled pools of carbamoyl phosphate and N-carbamoyl-aspartate, particularly in sgPYG cells, without appreciably changing their ^15^N-labeled fractions (**Figures 4B and S6B**). This was accompanied by accumulation of labeled and unlabeled CMP and reduced labeling of CTP, consistent with altered pyrimidine biosynthesis downstream of precursor formation. We next examined the contribution of glycogen-derived carbon using U-^13^C-glucose pulse-chase tracing. Disruption of either glycogenolytic pathway reduced labeling of N-carbamoyl-aspartate and eliminated detectable PRPP m+2, whereas unlabeled PRPP accumulated preferentially in sgGAA cells (**Figures 4C and S6C**). Labeled and unlabeled AMP also accumulated, most prominently following GAA deletion, accompanied by increased ATP abundance (**Figure 4C**). Thus, nitrogen and carbon tracing revealed distinct metabolic consequences of disrupting cytoplasmic and lysosomal glycogenolysis, suggesting that these pathways contribute differently to nucleotide biosynthesis.

**Figure 4.**
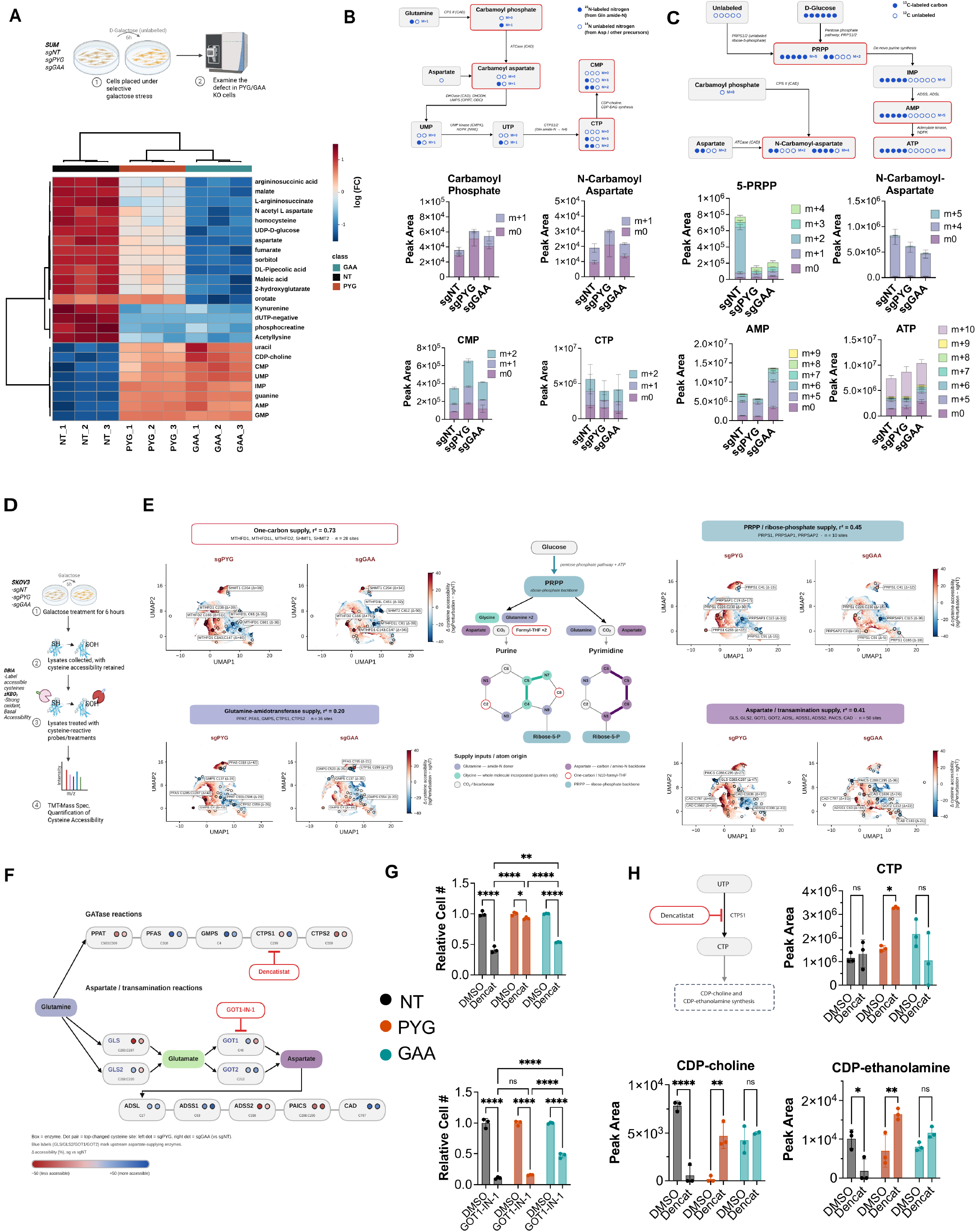
Cytoplasmic and lysosomal glycogenolysis differentially regulate nucleotide biosynthesis and metabolic dependencies. **(A)** Top altered polar metabolites in SUM159PT cells after 6 h in galactose (log fold change, row normalized, hierarchical clustering). **(B)** Amide-^15^N-glutamine tracing (12 h) in galactose. Top, incorporation of glutamine amide nitrogen into pyrimidine intermediates (filled circles, ^15^N). Bottom, isotopologues of carbamoyl phosphate, N-carbamoyl-aspartate, CMP, and CTP. **(C)** U-^13^C-glucose pulse-chase (24 h label, 6 h unlabeled galactose chase). Top, incorporation of glucose carbon into nucleotide precursors (filled circles, ^13^C). Bottom, isotopologues of 5-phosphoribosyl-1-pyrophosphate (PRPP), N-carbamoyl-aspartate, AMP, and ATP. **(D)** Cysteine accessibility proteomics. SKOV3 cells were transferred to galactose for 6 h, lysed under conditions that preserve native cysteine accessibility, treated with or without the oxidant KBO3 to define basal accessibility, labeled with desthiobiotin-iodoacetamide (DBIA), and analyzed by TMT-based mass spectrometry. **(E)** UMAP of quantified cysteine sites colored by change in reactivity relative to sgNT, in sgPYG (left of each pair) and sgGAA (right) cells, for four supply inputs into nucleotide biosynthesis: one-carbon (MTHFD1, MTHFD1L, MTHFD2, SHMT1, and SHMT2), PRPP/ribose phosphate (PRPS1, PRPSAP1, and PRPSAP2), aspartate/transamination (GLS, GLS2, GOT1, GOT2, ADSL, ADSS1, ADSS2, PAICS, and CAD), and glutamine amidotransferase (GATase; PPAT, PFAS, GMPS, CTPS1, and CTPS2). Sites in each category are circled, and the largest changes are labeled (Δ, percentage points). *r*^2^, sgPYG-sgGAA concordance within each category. Center, atom-level supply of purine and pyrimidine rings. **(F)** GATase and aspartate/transamination reactions annotated with the top-changed cysteine per enzyme (largest |ΔsgPYG| or |ΔsgGAA|). Left and right dots show the change in accessibility relative to sgNT in sgPYG and sgGAA cells, respectively. The CTPS1 inhibitor dencatistat and the GOT1 inhibitor GOT1-IN-1 are indicated. **(G)** Relative cell number of cells in galactose treated overnight with DMSO, 10 μM dencatistat (top), or 10 μM GOT1-IN-1 (bottom). **(H)** CTP, CDP-choline, and CDP-ethanolamine in cells treated with DMSO or 10 μM dencatistat for 24 h (schematic, left). Data are represented as mean ± SEM; n = 3 biological replicates; one-way ANOVA with multiple comparisons (\**p* < 0.05, \*\**p* < 0.01, \*\*\**p* < 0.001, \*\*\*\**p* < 0.0001; ns, not significant). sgNT, non-targeting control; sgPYG, *PYGB*/*PYGL* double knockout; sgGAA, *GAA* knockout. See also Figure S6.

To identify the metabolic pathways differentially affected by cytoplasmic and lysosomal glycogenolysis, we performed cysteine accessibility proteomics in sgNT, sgPYG, and sgGAA cells following 6 h of galactose exposure (**Figure 4D**). This approach identifies changes in pathway engagement through alterations in cysteine accessibility^42,43^, which can reflect changes in protein conformation and functional state^42,44^. We compared cysteine accessibility changes across four principal inputs into nucleotide biosynthesis: one-carbon metabolism, aspartate transamination, ribose phosphate supply, and glutamine amidotransferase reactions^13,45^. Changes in one-carbon metabolism were highly concordant between sgPYG and sgGAA cells (r^2^ = 0.725), whereas ribose phosphate supply (r^2^ = 0.45) and aspartate transamination (r^2^ = 0.41) showed weaker concordance. Glutamine amidotransferase reactions exhibited the greatest divergence (r^2^ = 0.20), suggesting that the two glycogenolytic pathways differentially influence enzymes involved in glutamine-dependent nucleotide biosynthesis (**Figure 4E**).

We focused on aspartate transamination and glutamine amidotransferase reactions, the two least concordant pathways, both of which are functionally linked to glutamine metabolism. Mapping cysteine accessibility changes onto these pathways identified distinct responses in several enzymes, including cytidine triphosphate synthase 1 (CTPS1) and glutamic-oxaloacetic transaminase 1 (GOT1) (**Figure 4F**). To determine whether these differences were associated with distinct metabolic dependencies, we treated cells with the CTPS1 inhibitor dencatistat ^46,47^ or the GOT1 inhibitor GOT1-IN-1^48^. Dencatistat reduced sgNT cell number by approximately 60% and sgGAA cells by approximately 50%, whereas sgPYG cells were largely unaffected (**Figure 4G**, top). Conversely, GOT1-IN-1 near-completely depleted sgNT and sgPYG cells, while sgGAA cells retained approximately 50% of their number (**Figure 4G**, bottom). In each case, cells were protected from inhibition of the enzyme whose top-ranked cysteine became less accessible in that background, and remained sensitive where accessibility increased (CTPS1 in sgPYG, GOT1 in sgGAA; **Figure 4F**). Because neither inhibitor is cysteine-directed, we interpret these accessibility changes as reporting reduced engagement of the corresponding reaction rather than altered inhibitor binding, consistent with the route of glycogen mobilization determining which of these reactions cells depend on.

To investigate the differential response to CTPS1 inhibition, we profiled nucleotide and phospholipid biosynthetic intermediates following dencatistat treatment. Despite their relative resistance to the inhibitor, sgPYG cells accumulated CTP and CDP-choline, with a smaller, nonsignificant increase in CDP-ethanolamine (**Figure 4H**). Dencatistat also increased glyceraldehyde-3-phosphate, dihydroxyacetone phosphate, and sn-glycerol-3-phosphate in sgPYG cells, whereas these metabolites were unchanged or reduced in sgGAA cells (**Figure S6D**). Notably, the increase in sn-glycerol-3-phosphate was specific to the sgPYG background, while UTP abundance remained largely unchanged. CDP-choline and CDP-ethanolamine are consumed together with diacylglycerol during phospholipid synthesis through the Kennedy pathway, with glycerol-3-phosphate contributing to the diacylglycerol backbone^49^. The concurrent accumulation of these intermediates in sgPYG cells is consistent with impaired utilization of CDP-alcohols downstream of their synthesis, potentially explaining why CTPS1 inhibition imposes a smaller additional growth defect in these cells. By contrast, sgGAA cells remained sensitive to dencatistat, indicating that loss of lysosomal glycogenolysis does not confer the same protection. We similarly examined the metabolic consequences of GOT1 inhibition. Treatment with GOT1-IN-1 preferentially reduced aspartate abundance in sgGAA cells, with comparatively smaller effects on α-ketoglutarate, glutamate, and oxaloacetate (**Figure S6E**). Thus, although both glycogenolytic pathways support mitochondrial metabolism and nucleotide biosynthesis, their disruption produces distinct changes in nucleotide precursor utilization and sensitivity to enzymes involved in glutamine-dependent metabolism. These findings suggest that cytoplasmic and lysosomal glycogenolysis make complementary contributions to nucleotide metabolism rather than functioning as redundant sources of glucose.

### The glycogen particle regulates pyrimidine biosynthesis and pyrimidinosome formation

Beyond supplying metabolic substrates, glycogen can regulate cellular signaling and metabolism through its spatial organization, including sequestration of Hippo pathway regulators and compartmentalization of pentose phosphate pathway enzymes^14,39^. To determine whether the glycogen particle similarly influences nucleotide biosynthesis, we compared cells lacking glycogen synthesis (sgGYS1) with those deficient in cytoplasmic (sgPYG) or lysosomal (sgGAA) glycogenolysis. Whereas disruption of glycogenolysis caused glycogen accumulation, GYS1 deletion depleted cellular glycogen (**Figures 5A and S7A**). Cysteine accessibility profiling revealed distinct changes in nucleotide biosynthetic enzymes between glycogen synthesis- and glycogenolysis-deficient cells (**Figure 5B**). We focused on pyrimidine biosynthesis, consistent with its association with the glycogenolysis signature in human tumors (**Figure 1B**) and the observation that uridine supplementation selectively rescued the growth defect of sgGYS1 cells under galactose stress (**Figure S7B**).

**Figure 5.**
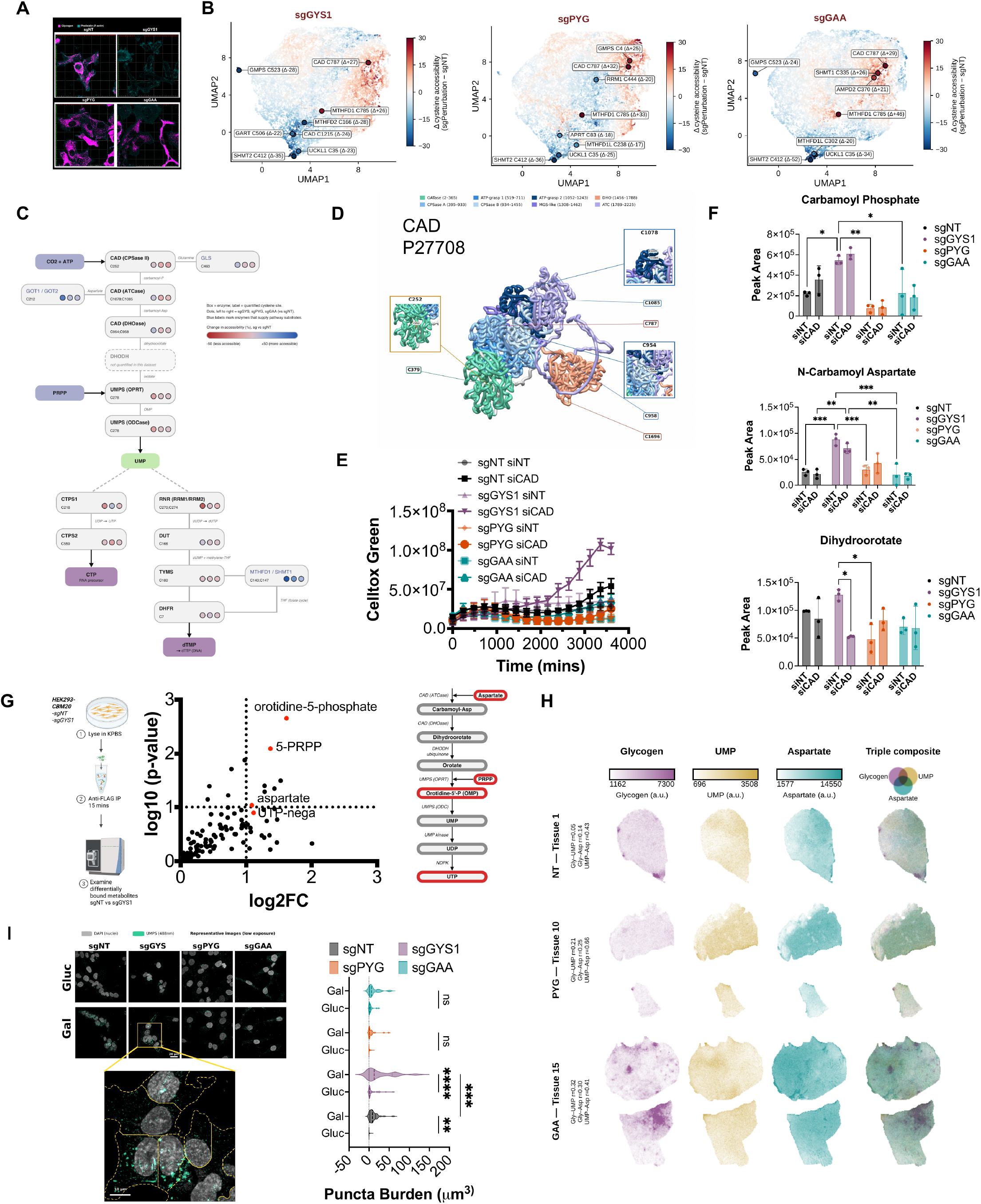
The glycogen particle spatially modulates pyrimidine biosynthesis and pyrimidinosome formation. **(A)** Representative immunofluorescence of glycogen (magenta) and F-actin (phalloidin, cyan) in SKOV3 sgNT, sgGYS1, sgPYG, and sgGAA cells. **(B)** UMAP of quantified cysteine sites colored by change in reactivity relative to sgNT in sgGYS1, sgPYG, and sgGAA cells. The largest-changing nucleotide metabolism sites are labeled. **(C)** De novo pyrimidine biosynthesis with one quantified cysteine mapped onto each enzyme: CAD (carbamoyl-phosphate synthetase II [CPSase II], aspartate transcarbamoylase [ATCase], and dihydroorotase [DHOase] domains), GLS, GOT1/GOT2, UMPS (OPRT and ODCase domains), CTPS1/CTPS2, RNR (RRM1/RRM2), DUT, TYMS, MTHFD1/SHMT1, and DHFR. Dots (left to right: sgGYS1, sgPYG, sgGAA) show the change in accessibility relative to sgNT; blue labels mark substrate-supplying enzymes. DHODH was not quantified (gray). **(D)** AlphaFold model of CAD (UniProt: P27708) showing the cysteines in (C) in the CPSase II (C252), ATCase (C1078 and C1085), and DHOase (C954 and C958) domains and additional sites (C379, C787, and C1696). Insets, C252, C1078, and C954 regions. **(E)** Cell death over time (CellTox Green fluorescence) of cells transfected with non-targeting (siNT) or *CAD* (siCAD) siRNA. **(F)** Carbamoyl phosphate, N-carbamoyl-aspartate, and dihydroorotate in siNT- and siCAD-transfected cells. **(G)** Rapid glycogen immunoprecipitation-metabolomics. HEK293 cells expressing FLAG-tagged CBM20 (sgNT or sgGYS1) were lysed in KPBS and subjected to a 15-min anti-FLAG immunoprecipitation, and bound metabolites were profiled by LC-MS (left). Volcano plot of metabolites enriched in sgNT relative to sgGYS1 pulldowns (middle; two-tailed t test; dashed lines, p = 0.1 and log_2_ fold change = 1). Enriched intermediates are outlined in red on the pyrimidine pathway (right). **(H)** MALDI-MSI of glycogen, UMP, and aspartate, and their composite, in representative sgNT (section 1), sgPYG (section 10), and sgGAA (section 15) SUM159 xenograft sections. Per-section pixel Spearman correlations are shown at left. **(I)** UMPS puncta. Left, representative low-exposure confocal projections of nuclei (DAPI, gray) and UMPS (green) in SKOV3 cells in glucose (Gluc) or galactose for 5 h (Gal); the inset shows the boxed sgGYS1 field with nucleus-seeded Voronoi cell territories (yellow dashed lines). Scale bars, 20 μm and 10 μm (inset). Right, UMPS puncta burden (total puncta volume per cell, μm^3^; n = 15–67 cells from 3–4 fields per condition). Puncta burden was compared by two-way ANOVA with Tukey’s multiple comparisons test. Data are represented as mean ± SEM; n = 3 biological replicates unless otherwise stated; one-way ANOVA with multiple comparisons, except (I), two-way ANOVA with Tukey’s multiple comparisons test (\**p* < 0.05, \*\**p* < 0.01, \*\*\**p* < 0.001, \*\*\*\**p* < 0.0001; ns, not significant). sgGYS1, *GYS1* knockout; sgNT, non-targeting control; sgPYG, *PYGB*/*PYGL* double knockout; sgGAA, *GAA* knockout. See also Figures S7 and S8.

Mapping cysteine accessibility changes onto the pyrimidine biosynthetic pathway identified alterations in several enzymes, including the multifunctional enzyme CAD, which catalyzes the first three steps of de novo pyrimidine biosynthesis (**Figure 5C**). Several cysteines within its carbamoyl phosphate synthetase II, aspartate transcarbamoylase, and dihydroorotase domains exhibited altered accessibility following glycogenolysis disruption, including C252, C1078/C1085, and C954/C958, respectively (**Figure 5D**). To determine whether glycogen availability alters dependence on CAD, we depleted CAD by siRNA in sgNT, sgGYS1, sgPYG, and sgGAA cells (**Figure S7C**). Strikingly, CAD depletion induced pronounced cell death specifically in sgGYS1 cells, whereas cells lacking either glycogenolytic pathway were comparatively resistant (**Figure 5E**). This selective vulnerability was accompanied by accumulation of carbamoyl phosphate and N-carbamoyl-aspartate and depletion of dihydroorotate in sgGYS1 cells following CAD silencing (**Figure 5F**). CAD depletion also significantly reduced UTP abundance in sgGYS1 cells, an effect not observed in the other genotypes (**Figure S7D**). Thus, loss of glycogen synthesis creates a distinct dependence on CAD-mediated pyrimidine biosynthesis, revealing a function for stored glycogen that differs from its metabolic utilization through glycogenolysis.

To investigate whether this function involves the association of pyrimidine biosynthetic intermediates with the glycogen particle, we developed a rapid glycogen immunoprecipitation approach using a FLAG-tagged carbohydrate-binding module (CBM20) to capture glycogen and its associated proteins and metabolites. Recovery of the established glycogen-interacting proteins PYGB, glycogen synthase, and GAA validated the approach (**Figure S7E**). Coupling rapid immunoprecipitation to metabolomic profiling identified several nucleotide biosynthetic intermediates enriched in glycogen-containing fractions relative to glycogen-deficient controls, including aspartate, 5-phosphoribosyl-1-pyrophosphate (PRPP), orotidine-5′-phosphate, and UTP (**Figures 5G and S7F**). Analysis of sgPYG and sgGAA cells further revealed increased recovery of orotidine-5′-phosphate, PRPP, and UTP under conditions of glycogen accumulation (**Figure S7G**). These findings indicate that metabolites required for pyrimidine biosynthesis associate with glycogen-containing structures, suggesting that stored glycogen influences their intracellular availability.

To examine this relationship in tumors, we analyzed the spatial distributions of glycogen, UMP, and aspartate using MALDI mass spectrometry imaging (**Figures 5H and S8A**). UMP and aspartate exhibited spatial correlation in control tumors, which decreased following disruption of cytoplasmic or lysosomal glycogenolysis, from a mean Spearman correlation of 0.50 in sgNT tumors to 0.33 and 0.24 in sgPYG and sgGAA tumors, respectively (**Figures 5H and S8B-S8C**). By contrast, spatial correlations between glycogen and either UMP or aspartate were generally weak, although individual glycogenolysis-deficient tumors exhibited increased colocalization (**Figure S8C**). These observations suggest that glycogen accumulation can alter the spatial relationships among pyrimidine biosynthetic metabolites, consistent with a role for the glycogen particle in their organization.

*De novo* nucleotide biosynthesis can involve the spatial assembly of enzymes and intermediates into multienzyme complexes, including purinosomes^50,51^ and pyrimidinosomes^52^ which depend on mitochondrial aspartate supply^53^. We therefore examined whether glycogen availability influences pyrimidinosome formation by measuring the accumulation of punctate UMP synthase (UMPS), a component of the pyrimidine biosynthetic machinery. Galactose exposure increased UMPS puncta in control cells, with a substantially greater increase following GYS1 deletion (**Figure 5I**). Conversely, disruption of either cytoplasmic or lysosomal glycogenolysis attenuated UMPS puncta formation (**Figure 5I**). These opposing effects suggest that stored glycogen influences the assembly or maintenance of pyrimidine biosynthetic complexes, potentially by regulating the availability of pathway intermediates. Together, these findings identify two complementary functions for glycogen in nucleotide metabolism: glycogenolysis supports the metabolic inputs required for nucleotide biosynthesis, whereas the glycogen particle influences the organization of pyrimidine biosynthetic intermediates and enzymes. The association of nucleotide precursors with glycogen-containing structures, together with the opposing effects of glycogen synthesis and degradation on UMPS puncta, supports a model in which glycogen availability regulates pyrimidine metabolism through spatial compartmentalization.

## DISCUSSION

Glycogen accumulation is a prominent feature of many cancers, yet its contribution to tumor metabolism remains incompletely understood. Here, we identify a role for glycogen in coordinating mitochondrial metabolism and nucleotide biosynthesis that extends beyond its conventional function as a glucose reserve. Glycogenolysis supports mitochondrial pyruvate oxidation, NADPH homeostasis, and reductive glutamine metabolism, with consequences for aspartate availability and nucleotide biosynthesis. Importantly, cytoplasmic and lysosomal glycogenolysis make distinct contributions to these processes, despite both pathways mobilizing stored glucose. We further find that the glycogen particle itself influences pyrimidine biosynthesis, with loss of glycogen synthesis creating a selective dependence on CAD and altering the organization of UMPS-containing structures. These findings establish complementary metabolic and spatial functions for glycogen in cancer cells.

A central finding is that glycogenolysis supports mitochondrial biosynthetic metabolism under nutrient stress. Glycogen breakdown has previously been linked to NADPH generation through the pentose phosphate pathway, particularly during hypoxia in cancer cells and in CD8⁺ memory T cells^8,39^. Our findings extend this relationship to mitochondrial carbon metabolism. Disruption of glycogenolysis impairs pyruvate oxidation in tumors and selectively reduces glutamine incorporation into proline and ornithine, while largely preserving oxidative glutamine labeling of TCA cycle intermediates. These changes are accompanied by impaired NADPH homeostasis and reduced aspartate availability, suggesting that glycogenolysis supports the reductive and biosynthetic functions of mitochondria. This is consistent with the emerging view that mitochondrial metabolism is functionally heterogeneous, with distinct metabolic pathways supporting ATP production and biosynthesis^26^. Glycogen may be particularly suited to supporting these processes because its cycle of synthesis mobilization provides carbon without requiring the reducing equivalents consumed during fatty acid synthesis. However, the mechanism linking glycogen turnover to reductive glutamine metabolism remains unresolved. Glycogen-dependent regulation of AMPK^54,55^, redox-sensitive glycogen phosphorylase activity^56^, and interactions between carbohydrate and lipid metabolism^57,58^ provide potential regulatory mechanisms. Whether these pathways directly govern glycogen-dependent mitochondrial metabolism, or whether the effects arise primarily through substrate availability and redox balance, will require further investigation.

Our findings also distinguish the metabolic functions of cytoplasmic and lysosomal glycogenolysis. Although disruption of either pathway impairs growth under nutrient stress, isotope tracing and cysteine accessibility profiling reveal distinct effects on nucleotide precursor metabolism. Cytoplasmic glycogenolysis is more closely associated with altered glutamine-dependent nitrogen utilization, whereas lysosomal glycogenolysis has a greater influence on specific carbon-derived nucleotide intermediates. These differences are reflected in the selective resistance of PYG-deficient cells to CTPS1 inhibition and of GAA-deficient cells to GOT1 inhibition. Thus, the two routes of glycogen breakdown appear to make complementary contributions to nucleotide biosynthesis rather than simply compensating for one another. How this specialization is established remains an important question. Glycogen metabolism is regulated by growth factor signaling, including the PI3K/AKT pathway, which controls glycogen synthesis and phosphorylase activity^7,59,60^. Compartment-specific signaling or differences in the intracellular availability of glycogen-derived metabolites may therefore contribute to the distinct metabolic consequences of cytoplasmic and lysosomal glycogenolysis. In particular, the relationship between lysosomal glycogen breakdown, aspartate transamination, and NADPH homeostasis warrants further investigation.

Perhaps the most unexpected finding is that the glycogen particle influences nucleotide biosynthesis independently of its degradation. Previous studies have established that glycogen can regulate signaling through the spatial sequestration of proteins, including components of the Hippo pathway in liver cancer^14^, and can associate with enzymes involved in pentose phosphate pathway metabolism^32^. Our findings extend this concept to pyrimidine biosynthesis. Glycogen synthesis-deficient cells exhibit a selective dependence on CAD, whereas glycogenolysis-deficient cells accumulate glycogen without developing the same vulnerability. Moreover, several pyrimidine biosynthetic intermediates associate with the glycogen particle, and glycogen availability alters the spatial distribution of UMP and aspartate and the formation of UMPS puncta. These observations suggest that stored glycogen influences the intracellular organization of pyrimidine metabolism. *De novo* nucleotide biosynthesis can be facilitated by the assembly of enzymes into multienzyme complexes, including purinosomes^50,51^ and pyrimidinosomes^52^, which depend on mitochondrial aspartate supply^53^. The opposing effects of glycogen synthesis and degradation on UMPS organization raise the possibility that the glycogen particle modulates the availability of intermediates for these biosynthetic complexes. Whether this reflects direct metabolite sequestration, altered enzyme assembly, or changes in local substrate availability remains to be established.

These findings suggest that glycogen abundance and glycogen turnover have distinct consequences for nucleotide metabolism. Glycogen degradation supports the mitochondrial production and utilization of biosynthetic precursors, whereas the stored glycogen particle may influence their accessibility to the nucleotide biosynthetic machinery. These roles may not function independently. During nutrient stress, glycogen mobilization could simultaneously support mitochondrial metabolism and alter the intracellular distribution of metabolites associated with the glycogen particle. Conversely, glycogen accumulation in the absence of degradation may alter nucleotide precursor availability without providing the metabolic inputs required for sustained biosynthesis. This distinction offers a potential explanation for why glycogen accumulation does not necessarily correlate with its contribution to tumor growth and highlights the importance of considering glycogen turnover, rather than glycogen abundance alone, when evaluating its metabolic function.

The dependence on glycogenolysis is particularly evident under conditions that restrict glucose utilization, suggesting that glycogen metabolism provides an adaptive mechanism for maintaining biosynthetic activity during nutrient limitation. The impaired growth of glycogenolysis-deficient tumors further supports its relevance to cancer metabolism *in vivo*. Whether the metabolic and spatial functions described here are conserved across different tumor environments, and whether they create therapeutic vulnerabilities that can be exploited selectively, remain important questions. In particular, the distinct sensitivities to CTPS1 and GOT1 inhibition suggest that the route of glycogen mobilization may influence dependence on individual nucleotide biosynthetic enzymes. More broadly, our findings establish glycogen as both a metabolic reserve and a regulator of biosynthetic organization, linking the mobilization of stored carbon to mitochondrial metabolism and the spatial control of nucleotide biosynthesis.

## LIMITATIONS OF THE STUDY

Although our findings establish distinct metabolic functions for cytoplasmic and lysosomal glycogenolysis, the mechanisms underlying their differential contributions to nucleotide biosynthesis remain unresolved, particularly their relationships with CTPS1 and GOT1. The extent to which glycogen structure and posttranslational modifications influence these processes was not examined, and tools to selectively or simultaneously perturb the two glycogenolytic pathways remain limited. Finally, although our findings implicate the glycogen particle in the spatial organization of pyrimidine metabolism, whether glycogen directly sequesters biosynthetic intermediates or regulates metabolic channeling remains to be determined.

## RESOURCE AVAILABILITY

### Lead Contact

Further information and requests for resources and reagents should be directed to and will be fulfilled by the Lead Contact, Alex Toker.

### Materials Availability

All unique reagents generated in this study are available from the lead contact, Alex Toker, upon request.

### Data and Code Availability

All raw data (original western blot images and other raw data files) and original code have been deposited to the Harvard Dataverse (https://doi.org/10.7910/DVN/AQLHPQ) and will be made available as of the date of publication.

Any additional information will be made available from the lead contact upon request.

## ACKNOWLEDGMENTS

T.K. is supported by the American Cancer Society (PF-25-1379136-01-PFMBB). S.A.H. is supported by the M1/IAM 2003 Rankin-Rasmussen Postdoctoral Fellowship (PF-23-1018904-01-ET) from the American Cancer Society. M.C.C. is supported by an NIH F32 NRSA Fellowship (F32DK149569) and a T32 Molecular Imaging in Cancer Biology (MICB) Research Fellowship (T32CA254875). J.M.A. is supported by NIH grant P01CA120964. M.S.G. is supported by NIH grant R35NS116824. M.S.G. and R.C.S. are supported by NIH grants R01AG096366, R01CA266004, R01CA288696, and R01AG078702. K.R.K. is supported by NIH grants R01CA237466, R01CA252037, R01CA248364, R01CA249294, and R01CA283578, and by the Center for Molecular Imaging and Bioengineering (CMIB) and the Experimental Therapeutics Center at MSKCC. L.B.-P. is supported by the Ludwig Center at Harvard, the Mark Foundation for Cancer Research, the American Association for Cancer Research (19-20-45-BARP), the American Cancer Society, the Melanoma Research Alliance, Lungevity, ALK Positive, the V Foundation, the Mary Kay Foundation, the Paula and Rodger Riney Foundation, Mark and Lisa Shwartz, the Pew-Stewart Trusts, and the NIH/NCI (R01CA285415, R37CA260062). A.T. is supported by NIH grant CA253097 and the Ludwig Center at Harvard. This work was also supported by NIH/NCI Cancer Center Support Grants P30CA008748 (Memorial Sloan Kettering Cancer Center) and P30CA247796 (UF Health Cancer Institute).

We would like to thank the BIDMC Flow Cytometry (RRID:SCR_012305), Mass Spectrometry (RRID:SCR_017735), Preclinical Murine Pharmacogenetics (RRID:SCR_026599), Immunostaining and Microscopy Cores (RRID:SCR_012312) for facilitating this study. We would like to thank Professor Navdeep Chandel for providing AOX, LBNOX, and mitoLBNOX vectors used in this study, and Professor Dawang Zhou for providing CBM20 vectors used in this study. We are grateful to members of the Toker and Dibble laboratories for helpful discussions and critical feedback.

## AUTHOR CONTRIBUTIONS

Conceptualization: T.K., A.T.

Methodology: T.K., A.T., J.M.A., S.A.H., K.R.K., M.M.C., R.C.S., M.S.G., L.B.P, M.G., R.L.

Investigation: T.K., S.A.H., M.M.C., M.C.C., T.P., J.M.A., J.L., H. D., J.G.C. Formal Analysis: T.K., S.A.H., M.M.C., M.C.C., T.P., J.M.A.

Writing - Original Draft: T.K.

Writing - Review & Editing: All authors Supervision: A.T., T.K.

Funding Acquisition. A.T., T.K, M.M.C, K.R.K, M.C.C.

## SUPPLEMENTAL INFORMATION TITLES

Figure S1. Loss of cytoplasmic glycogenolysis restrains tumor growth and TCA cycle metabolite abundance in vivo, related to Figure 1

Figure S2. Cytoplasmic and lysosomal glycogenolysis are required for growth under galactose stress across cancer cell lines, related to Figure 2

Figure S3. Glycogen is mobilized within 1 h of galactose stress, and glycogen phosphorylase inhibition phenocopies genetic loss of glycogenolysis in TCA cycle labeling, related to Figure 2

Figure S4. Glycogenolysis loss alters pyruvate carbon fate and the malate/citrate labeling ratio and blunts rescue by NAD(P)H reoxidation, related to Figures 2 and 3

Figure S5. Glycogenolysis loss raises the NADP^+^/NADPH ratio and impairs reductive glutamine metabolism in vitro and in vivo, related to Figure 3

Figure S6. Glycogenolysis loss perturbs nucleotide pools and labeling and differentially alters sensitivity to CTPS1 and GOT1 inhibition, related to Figure 4

Figure S7. Glycogen synthesis loss creates a distinct pyrimidine dependency, and CBM20 glycogen immunoprecipitation captures glycogen-associated proteins and pyrimidine intermediates, related to Figure 5

Figure S8. Spatial co-distribution of glycogen with UMP and aspartate in xenograft tumors, related to Figure 5

## STAR★METHODS

### KEY RESOURCES TABLE

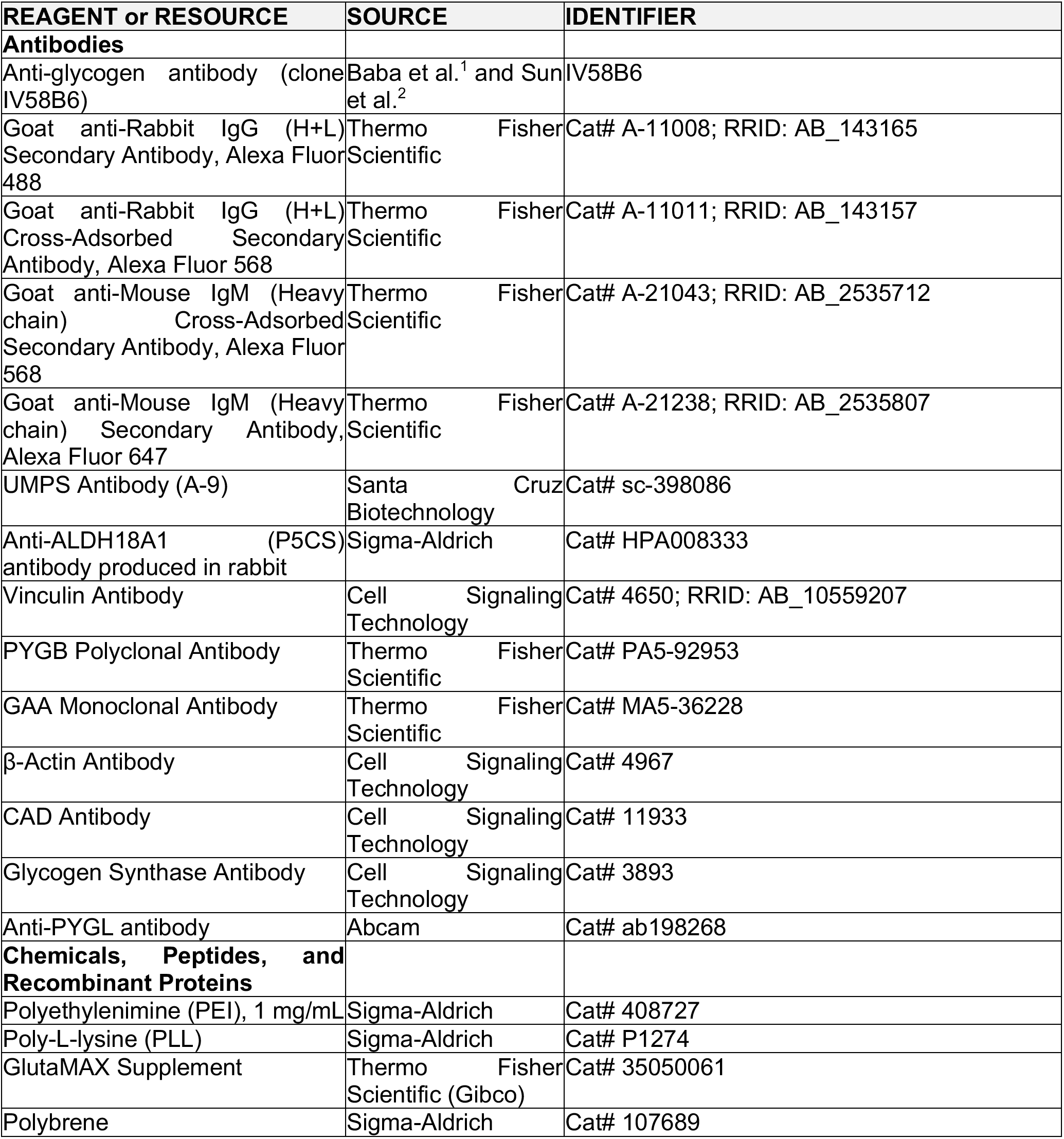
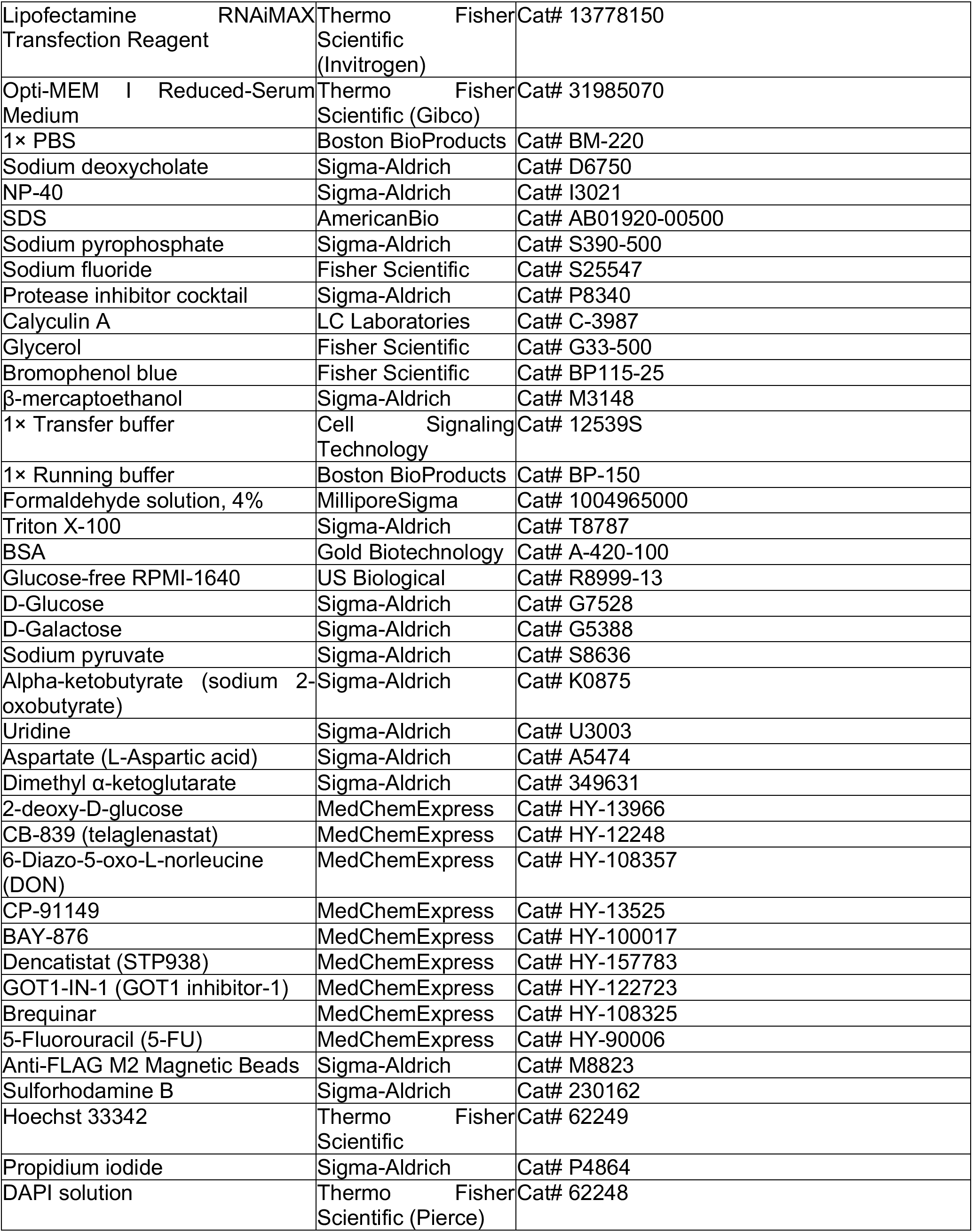
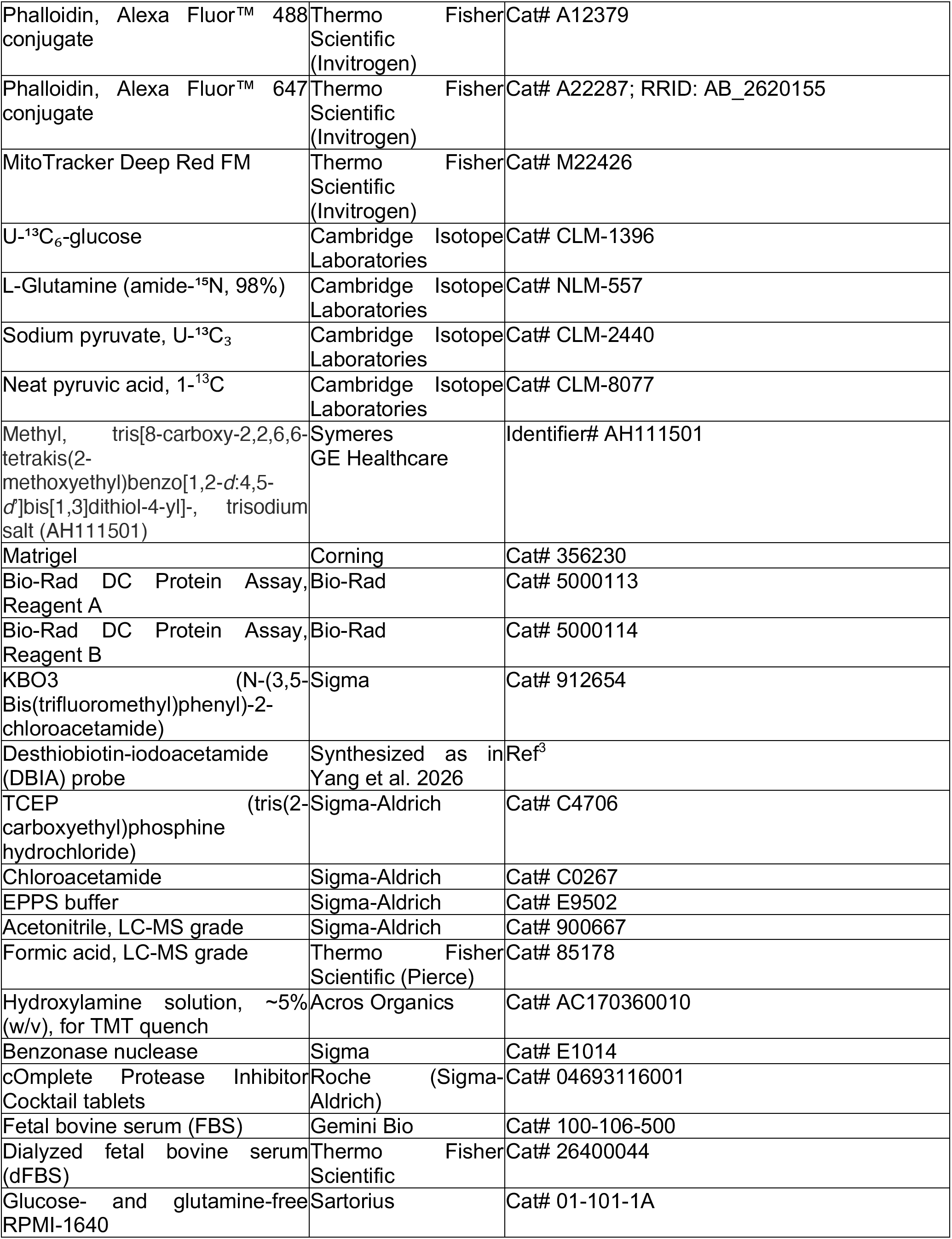
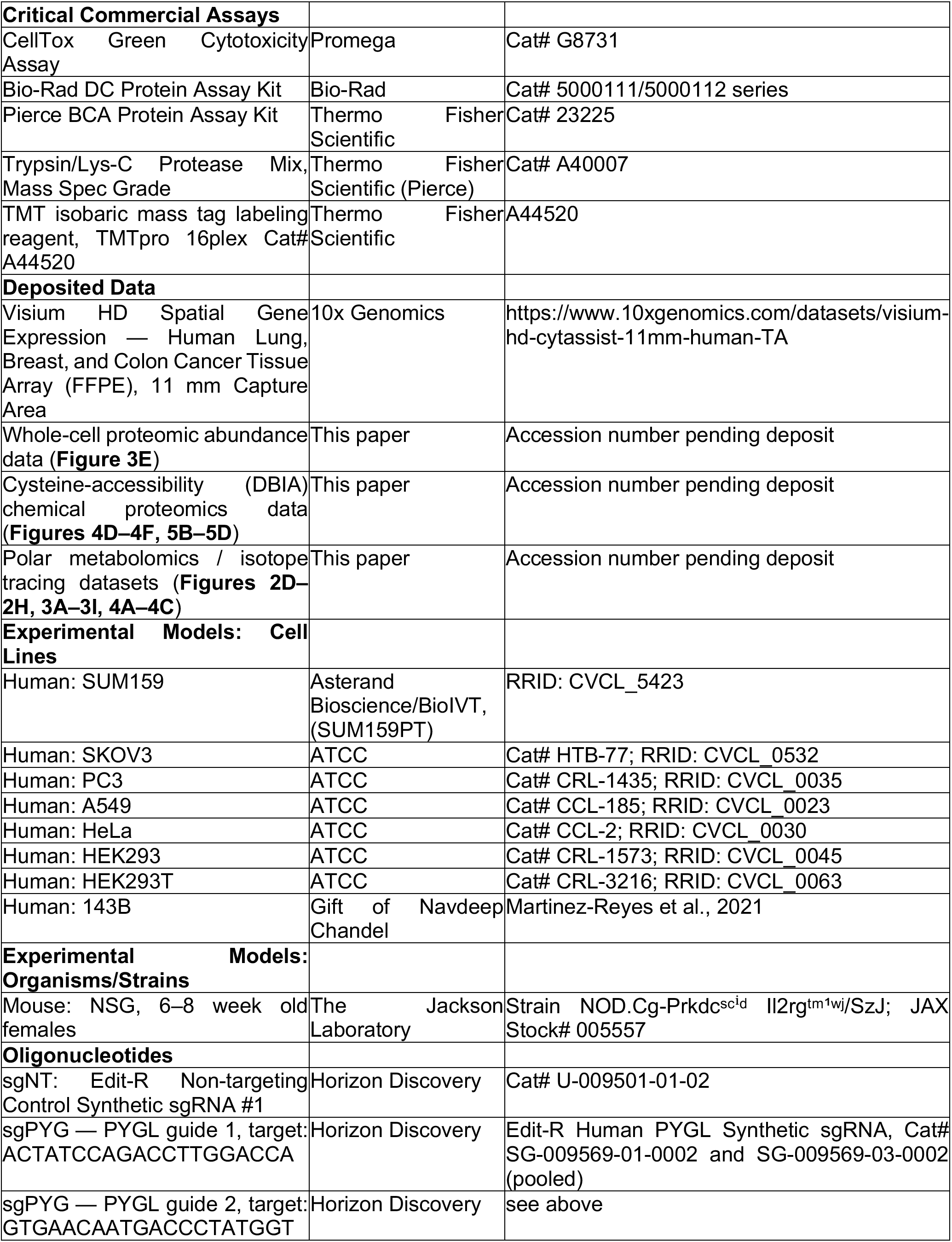
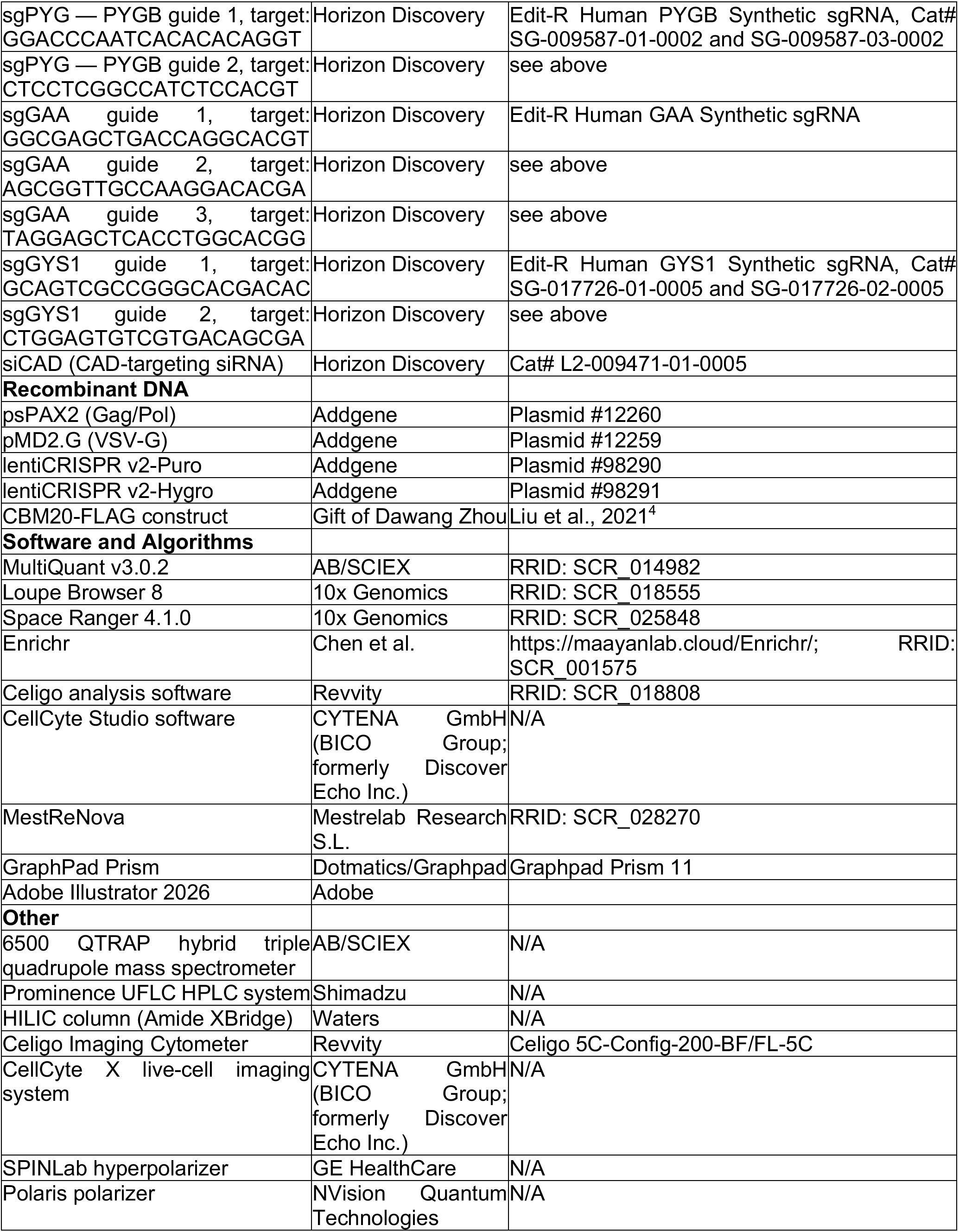
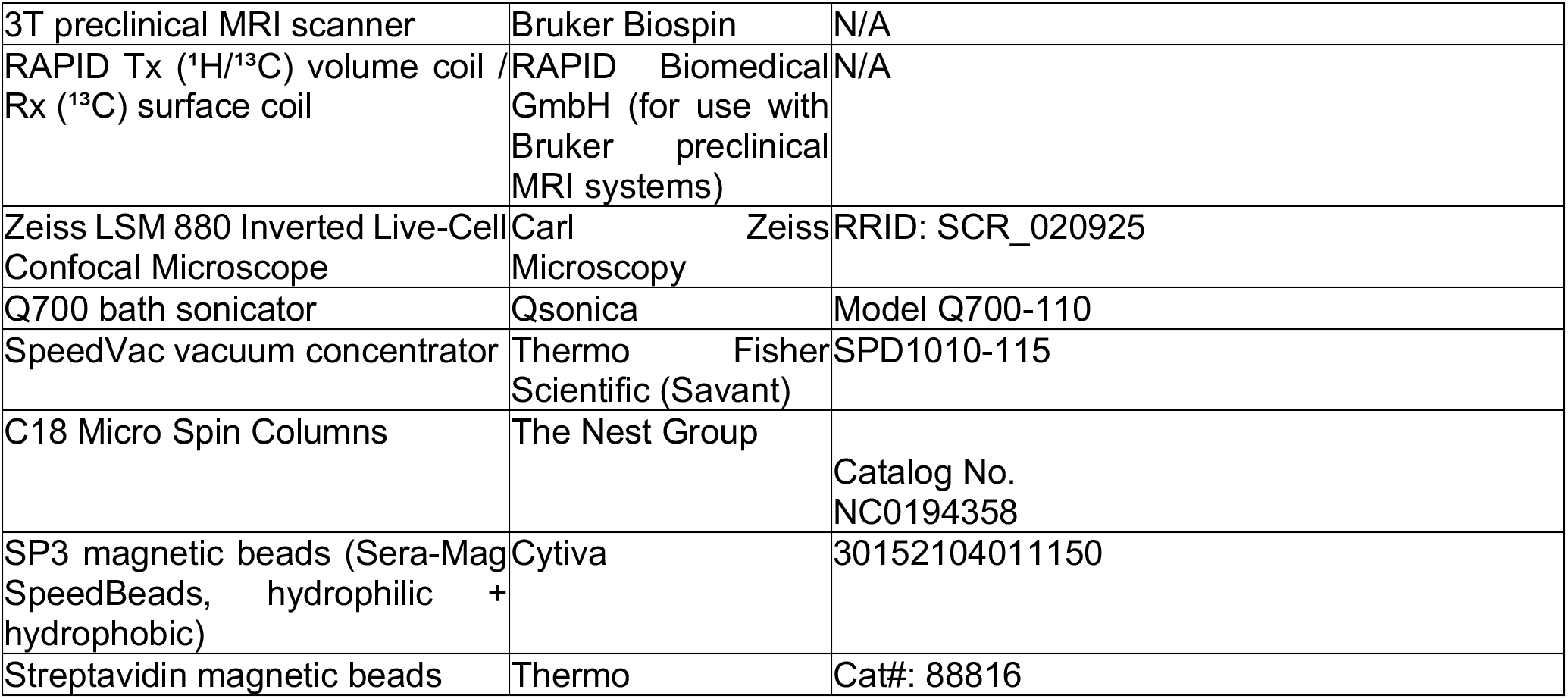

### EXPERIMENTAL MODEL AND STUDY PARTICIPANT DETAILS

#### Mice

Growth experiments were performed at Beth Israel Deaconess Medical Center and were approved by and conducted in accordance with the guidelines of the Beth Israel Deaconess Medical Center Institutional Animal Care and Use Committee. MRI experiments were performed at Memorial Sloan Kettering Cancer Center and were reviewed and approved by the Institutional Animal Care and Use Committee at Memorial Sloan Kettering Cancer Center, in accordance with protocol number 13-12-019. In both instances, 6-8 week old female NSG mice were purchased from Jackson Laboratories

#### Cell Lines

Cell lines were maintained at 37°C and 5% CO₂. SKOV3, PC3, A549, HeLa, HEK293T and HEK293 cells were obtained from ATCC. SUM159 cells (SUM159PT) were from Asterand Bioscience/BioIVT, 143B cells were a gift from Navdeep Chandel. HEK 293T, HEK293 and HeLa cells were cultured in DMEM with 10% fetal bovine serum (FBS; Gemini); all other lines were cultured in RPMI-1640 with 10% FBS. For assays, cells were seeded in base media with 10% dialyzed fetal bovine serum (dFBS; Thermo Fisher Scientific), and for carbon-source experiments, cells were plated in glucose-free media (US Biological, R8999-13) supplemented with 10 mM D-glucose or D-galactose (Sigma-Aldrich).

### METHOD DETAILS

#### Lentivirus Generation

Lentivirus was generated using a master mix of 11.1 µg psPAX2 (Gag/Pol), 0.6 µg VSV-G, 6.3 µg of plasmid DNA, and 1,200 µL of serum-free, antibiotic-free DMEM. Then, 54 µL of 1 mg/mL polyethylenimine (PEI; Sigma-Aldrich, 408727) was added, and the master mix was vortexed for 10 seconds and incubated at room temperature for 15 minutes. Tissue culture–treated 10-cm dishes were coated with poly-L-lysine (PLL; Sigma-Aldrich, P1274) for 5 minutes and then washed three times with 1× PBS. HEK293T cells were trypsinized and counted, and 9 × 10⁶ cells were resuspended in 7.8 mL of DMEM supplemented with 10% FBS. Approximately 1,200 µL of the reaction mixture was then added to the HEK293T cell suspension, mixed, and plated on the PLL-coated 10-cm dishes. GlutaMAX Supplement (Thermo Fisher Scientific, 35050061) was added to a final concentration of 1×. Cells were incubated for 48 hours. Virus was harvested from HEK293T cells by collecting media and passing it through a 10-mL syringe fitted with a 0.45-µm filter. Virus was either frozen at −80°C or used immediately to transduce target cells.

Transductions were performed in target cells seeded in 6-well plates at 200,000 cells/well. Thirty minutes prior to transduction, Polybrene (Sigma-Aldrich, 107689) was added at a final concentration of 5 µg/mL. Virus was added and cells were incubated at 37°C for 48 hours. The next day, media was changed to fresh media for 24 hours, and the following day, the relevant antibiotic was added; cells were maintained in antibiotic-containing media until non-transduced control cells had died.

#### Generation of Genetically Edited Cell Lines

For CRISPR-Cas9 editing, cells were first stably transduced with lentiCRISPR v2-Puro (Addgene, Plasmid #98290) or lentiCRISPR v2-Hygro (Addgene, Plasmid #98291).

Stably transduced cells were transfected with the relevant guide RNA using Lipofectamine RNAiMAX Transfection Reagent (Thermo Fisher Scientific, 13778150) as follows. All reagents were brought to room temperature before proceeding. Lipofectamine RNAiMAX was diluted in Opti-MEM reduced-serum media (Thermo Fisher Scientific, 31985070) at 1.5 µL Lipofectamine RNAiMAX per 25 µL Opti-MEM, vortexed, and incubated at room temperature for 5 minutes. sgRNAs were diluted from 10 µM stocks to a 2 µM guide transfection complex by adding 2 µL sgRNA to 8 µL Tris buffer in Opti-MEM media, vortexed, and incubated at room temperature for 5 minutes. For simultaneous use of multiple guides, 2 µM of each guide was used, and an equivalent amount of sgNT was used. Equal volumes of the Lipofectamine RNAiMAX/Opti-MEM mixture and the sgRNA/Opti-MEM mixture were then combined, vortexed, and incubated for 20 minutes at room temperature. For a lipofectamine-only control, equal volumes of Lipofectamine RNAiMAX/Opti-MEM and Opti-MEM alone were combined, vortexed, and incubated at room temperature for 20 minutes. 200 µL of the resulting mixture was added to each well.

To achieve maximal editing efficiency, transfected cells were single-cell cloned in 96-well plates. Clones were expanded, and successful gene editing was assessed by immunoblotting. Successfully edited clones were then pooled together (≥ 5 clones per pool) to generate each knockout cell line.

##### sgRNA target sequences

sgNT: Edit-R synthetic sgRNA non-targeting control (Horizon Discovery) U-009501-01-02, Sequence Unspecified

sgPYG: Edit-R CRISPR (knockout) Human PYGL Synthetic sgRNA SG-009569-01-0002 and SG-009569-03-0002 and Edit-R CRISPR (knockout) Human PYGB Synthetic sgRNA SG-009587-01-0002 and SG-009587-03-0002

PYGL 1 Target Sequence: ACTATCCAGACCTTGGACCA

PYGL 2 Target Sequence: GTGAACAATGACCCTATGGT

PYGB 1 SG-009587-01-0002 Target Sequence: GGACCCAATCACACACAGGT

PYGB 2 SG-009587-03-0002 Target Sequence: CTCCTCGGCCATCTCCACGT

sgGAA; Edit-R CRISPR (knockout) Human GAA Synthetic sgRNA (Horizon Discovery) SG-008881-01-0002

SG-008881-01 Target sequence: GGCGAGCTGACCAGGCACGT

SG-008881-02 Target sequence: AGCGGTTGCCAAGGACACGA

SG-008881-03 Target sequence: TAGGAGCTCACCTGGCACGG

sgGYS1: Edit-R CRISPR (knockout) Human GYS1 Synthetic sgRNA SG-017726-01-0005 and SG-017726-02-0005

SG-017726-01-0005 target sequence: GCAGTCGCCGGGCACGACAC

SG-017726-02-0005 target sequence: CTGGAGTGTCGTGACAGCGA

#### Immunoblotting

Cells were washed once in cold 1× PBS (Boston BioProducts, BM-220) and collected on wet ice in 4°C RIPA lysis buffer [150 mmol/L Tris-HCl, 150 mmol/L NaCl, 0.5% (w/v) sodium deoxycholate (Sigma-Aldrich, D6750), and 1% (v/v) NP-40 (Sigma-Aldrich, I3021); pH 7.5] containing 0.1% SDS (AmericanBio, AB01920-00500), 1 mmol/L sodium pyrophosphate (Sigma-Aldrich, S390-500), 20 mmol/L sodium fluoride (Fisher Scientific, S25547), 0.5% (v/v) protease inhibitor cocktail (104 mmol/L AEBSF, 80 µmol/L aprotinin, 4 mmol/L bestatin, 1.4 mmol/L E-64, 2 mmol/L leupeptin, and 1.5 mmol/L pepstatin A; Sigma-Aldrich, P8340), and 50 nmol/L calyculin A (LC Laboratories, C-3987), all added immediately before use. Plates were scraped into 1.5-mL microcentrifuge tubes, vortexed, and incubated on wet ice for 15 minutes, then centrifuged at 15,000 rpm for 10 minutes at 4°C. Supernatants were transferred to new tubes and protein concentrations were measured using the Bio-Rad DC Protein Assay (Bio-Rad; Reagent A, 5000113; Reagent B, 5000114). Sample concentrations were normalized using 2× SDS sample buffer [62.5 mmol/L Tris (pH 6.8), 2% SDS, 10% glycerol (Fisher Scientific, G33-500), bromophenol blue (Fisher Scientific, BP115-25), and 5% (v/v) β-mercaptoethanol (Sigma-Aldrich, M3148)]. Cell lysates were boiled at 95°C for 5 minutes and stored at −20°C.

Cell lysates were separated by SDS-PAGE on 7.6%–15% acrylamide gels in 1× running buffer (Boston BioProducts, BP-150), or on precast 4%–16% gradient gels (Thermo Fisher Scientific). For the former, proteins were transferred to nitrocellulose membranes at 100 V for 90 minutes in cold 1× transfer buffer (Cell Signaling Technology, 12539S); for the latter, proteins were transferred using the iBlot2 system (Thermo Fisher Scientific).

#### Immunofluorescence

Black-walled, glass-bottom, tissue culture–treated plates were coated with PLL (Sigma-Aldrich, P1274) for 5 minutes and then washed three times with 1× PBS. Cells were seeded at 100,000 cells/well in 80 µL. Cells were fixed with 4% formaldehyde solution (MilliporeSigma, 1004965000) for 1 hour at room temperature and then washed three times with 1× PBS. Cells were permeabilized with 0.01% Triton X-100 (Sigma-Aldrich) in PBS for 3 minutes, washed three times with 1× PBS, and blocked with 0.5% BSA (Gold Biotechnology, A-420-100) in 1× PBS for 1 hour, rocking at room temperature.

After blocking, cells were incubated with primary antibody — anti-glycogen antibody (clone IV58B6) — overnight at 4°C. The next day, plates were washed three times with 1× PBS and then incubated in 0.5% BSA with secondary antibody at 1:5000 dilution for 60 minutes, rocking at room temperature – either Goat anti-Rabbit IgG (H+L) Secondary Antibody, Alexa Fluor™ 488 (Thermo Fisher, A-11008) or Goat anti-Rabbit IgG (H+L) Cross-Adsorbed Secondary Antibody, Alexa Fluor™ 568 (Thermo Fisher, A-11011) for rabbit primary antibodies, or Goat anti-Mouse IgM (Heavy chain) Cross-Adsorbed Secondary Antibody, Alexa Fluor™ 568 (Thermo Fisher,A-21043) or Goat anti-Mouse IgM (Heavy chain) Secondary Antibody, Alexa Fluor™ 647 (Thermo Fisher, A-21238) for mouse IgM primary antibodies, including anti-glycogen. Where specified, DAPI solution (100 ng/ml, Thermo Fisher 62248) and Phalloidin (1:200, Thermo Fisher A12379 or A22287) were added to secondary antibody solution as counterstains.

Cells were then washed twice with 1× PBS and imaged using the Zeiss LSM 880 Inverted Live-Cell Confocal Microscope.

#### Galactose/Glucose-Free Culture Experiments

For carbon-source experiments in glucose-free, galactose, or glutamine-free media, cells were seeded overnight in 96-well plates in RPMI-1640 with 10% dialyzed FBS. The following day, cells were washed twice in warm 1× PBS and media was changed to the relevant assay media: for glucose, galactose, and glucose-free conditions, glucose-free RPMI-1640 supplemented with 10 mM glucose, galactose, or an isovolumetric amount of water; for glutamine experiments, glucose- and glutamine-free RPMI-1640 supplemented with 2 mM L-glutamine and either 10 mM glucose or 10 mM galactose. Where relevant, rescue agents were added 1 hour after the media change: 1 mM pyruvate (Sigma-Aldrich; 2 or 10 mM in Figure 2F), 1 mM alpha-ketobutyrate (Sigma-Aldrich), 100 µM uridine (Sigma-Aldrich), 1 mM aspartate (Sigma-Aldrich), 1 mM dimethyl α-ketoglutarate (Sigma-Aldrich, 349631), 1× MEM non-essential amino acids (NEAA; Thermo Fisher Scientific, 11140050), 1× GlutaMAX (Thermo Fisher Scientific, 35050061), 200 µM L-serine (Sigma-Aldrich, S4500), and 0.6 mM 2-deoxy-D-glucose (MedChemExpress). Cells were harvested at indicated timepoints for the relevant assays (TCA fixation for SRB growth assays; Hoechst/PI staining for cytotoxicity assays).

#### Drug Treatment Assays

Cells were seeded overnight in 96-well plates. The following day, they were treated with drugs at indicated concentrations: CB-839, 6-diazo-5-oxo-L-norleucine (DON), CP91149, BAY-876, dencatistat, GOT1-IN-1, brequinar, and 5-FU (all from MedChemExpress). Assays were ended at indicated timepoints by either TCA fixation or Hoechst/PI staining.

#### Glycogen Immunoprecipitation

Glycogen immunoprecipitation was performed in HEK293 cells stably expressing the CBM20-FLAG construct, using a modified version of the Lyso-IP protocol^5^. Briefly, cells were seeded in 15-cm plates overnight in RPMI with 10% dialyzed FBS. The following day, media was changed to fresh media and cells were allowed to equilibrate for 1 hour. Cells were then washed twice in cold PBS and scraped in 1 mL KPBS containing protease and phosphatase inhibitor, and collected in 2-mL Eppendorf tubes. Lysates were homogenized by passing 8 times through a 26-gauge syringe, after which a 100-µL sample was taken and mixed 1:1 with lysis buffer for downstream protein normalization. Lysates were incubated with 100 µL of pre-washed anti-FLAG M2 magnetic beads (Sigma-Aldrich) in the cold room, with rotation. Beads were then rapidly washed 3× for 5 minutes each in KPBS, and metabolites were eluted from the beads using 100% methanol for 5 minutes, after which the eluate was diluted to 80% (v/v) methanol using HPLC-grade H₂O and transferred to a new tube. Insoluble material was removed by centrifugation at 15,000 × g for 10 minutes at 4°C, after which the metabolite-containing supernatant was dried under nitrogen gas as described above.

#### Cell Density (Sulforhodamine B) Assays

Cells were seeded in tissue culture–treated 96-well plates in 80 to 100 µL of appropriate growth media (3,000 cells/well) and incubated for 24 hours; cell density was then assayed by sulforhodamine B (SRB; Sigma-Aldrich, 230162) staining, as previously described^6^, to determine the number of cells at the start of the experiment (day 0). Cells were then treated with 5 to 20 µL of drug for the indicated period, bringing the final volume in each well to 100 µL. At the endpoint, relative cell density was assayed by SRB staining. Cell density at each time point was normalized to the day-0 control. Unless otherwise stated in the figure legend, SRB assays were performed with n = 3 biological replicates; multi-day growth curves (Figure S2B and Figure 2C) were performed with n = 3 replicate wells per condition.

#### Cell Density (CellCyte Live-Cell Imaging) Assays

For temporal dynamics of long-term growth, cells were imaged using the CellCyte X live-cell imaging system (Discover Echo). Cells were seeded in 96-well plates and allowed to attach overnight. The next day, cells were placed into the CellCyte imaging system and images were taken using the 10X objective (2 images per well) every 4 hours for 6 days. Cell confluence was calculated using CellCyte software with the following settings: contrast sensitivity 50%, smoothing 2 a.u., filled hole size 100 µm^2^, min object size 100 µm^2^. Data was plotted as mean ± SEM.

#### Cytotoxicity Assays

Briefly, cells were treated as indicated and then stained for 15 minutes with 3 µg/mL Hoechst 33342 (Thermo Fisher Scientific) and 3 µg/mL propidium iodide (PI; Sigma-Aldrich). Cytotoxicity assays were performed on the Celigo Imaging Cytometer (Revvity). Briefly, plates were imaged using the two-color cell-viability setting (Dead+Total; Blue/DAPI and Red/RFP filters), and images were quantified using Celigo software to determine live cells (Hoechst-positive), dead cells (PI-positive), death % (PI-positive/Hoechst-positive × 100), and living cells (Hoechst-positive − PI-positive). Cell analysis settings were as follows: total (algorithm fluorescence, intensity threshold 2, precision high, cell diameter 5 pixels) and dead (algorithm fluorescence, intensity threshold 4, precision high, cell diameter 4 pixels).

#### Isotope Tracing and Steady-State Metabolomics Measurement

##### Cell seeding, drug treatments, and metabolite harvest

Cells were plated at 150,000 cells/mL in tissue culture–treated 6-cm dishes in the relevant base media with 10% dialyzed FBS. Three biological replicates were seeded for metabolomics analysis and two for protein analysis by immunoblotting. After treatment, plates were placed on wet ice and media were aspirated completely. Plates were washed once with 2 mL of cold PBS, aspirated completely, and moved to dry ice. On dry ice, 1.5 mL of −80°C 80% MeOH was added to each plate. Plates were scraped on wet ice, and 1.5 mL of each sample was transferred to a 2-mL Eppendorf tube and vortexed for 30 seconds. Samples were centrifuged at 20,000 × g at 4°C for 5 minutes and then immediately dried under nitrogen gas. Dried samples were stored at −80°C. To prepare samples for LC-MS analysis, dried samples were resuspended in 20 µL of mass spectrometry–grade water, centrifuged at 10,000 × g at 4°C for 3 minutes, and supernatants were transferred to mass spectrometry–compatible vials.

##### Heavy isotope labeling

Heavy isotope labeling experiments used U-¹³C-glucose, ¹⁵N-glutamine, and U-¹³C-pyruvate (Cambridge Isotope Laboratories). Cells were washed twice in warm PBS and labeled for the indicated times with the indicated tracers (10 mM glucose, 1 mM glutamine, or 1 mM pyruvate) in media lacking endogenous levels of the relevant metabolite — glucose-free media (US Biological), glucose- and glutamine-free media (Sartorius) supplemented with glucose, and RPMI-1640 without exogenous pyruvate, respectively. For U-¹³C-glucose experiments, cells were pulsed for 24 hours, followed by a chase for the indicated times in unlabeled media. For glutamine tracing, cells were labeled for 12 hours; for pyruvate rescue experiments, cells were labeled for 12 hours. Cells were harvested in 80% methanol as described above and processed for LC-MS as described below. For heavy-isotope-labeled metabolite tracing, SRMs were adapted from previous studies, with optimization of collision energies and, where needed, selection of the most reliable Q3 product ion ^7^

##### Targeted Metabolomics

Samples were re-suspended using 20 uL HPLC grade water for mass spectrometry. 5-7 μL were injected and analyzed using a hybrid 6500 QTRAP triple quadrupole mass spectrometer (SCIEX) coupled to a Prominence UFLC HPLC system (Shimadzu) via selected reaction monitoring (SRM) of a total of 300 endogenous water soluble metabolites for steady-state analyses of samples. Some metabolites were targeted in both positive and negative ion mode for a total of 311 SRM transitions using positive/negative ion polarity switching. ESI voltage was +4950V in positive ion mode and –4500V in negative ion mode. The dwell time was 3 ms per SRM transition and the total cycle time was 1.55 seconds. Approximately 9-12 data points were acquired per detected metabolite. Samples were delivered to the mass spectrometer via hydrophilic interaction chromatography (HILIC) using a 4.6 mm i.d x 10 cm Amide XBridge column (Waters) at 400 uL/min. Gradients were run starting from 85% buffer B (HPLC grade acetonitrile) to 42% B from 0-5 minutes; 42% B to 0% B from 5-16 minutes; 0% B was held from 16-24 minutes; 0% B to 85% B from 24-25 minutes; 85% B was held for 7 minutes to re-equilibrate the column. Buffer A was comprised of 20 mM ammonium hydroxide/20 mM ammonium acetate (pH=9.0) in 95:5 water:acetonitrile. Peak areas from the total ion current for each metabolite SRM transition were integrated using MultiQuant v3.0.2 software (SCIEX). NAD^+^/NADH and NADP^+^/NADPH ratios (Figure 3D and Figure S5A and S5B) were calculated as the ratio of the integrated peak areas of the oxidized and reduced cofactors from steady-state polar metabolomics of cells cultured in galactose for 6 h.

#### Cysteine-Focused Chemical Proteomics

Iso-TMT samples were prepared as previously described^3,8,9^. In brief, after relevant treatments, cells were pelleted, washed twice with ice-cold PBS, snap-frozen in liquid nitrogen, and stored at -80°C until use (n=3 biological replicates). Frozen cell pellets were lysed in DPBS supplemented with Benzonase (Sigma-Aldrich) and protease inhibitors (Roche) using a chilled bath sonicator (Q700, QSonica) and centrifuged for 3 min at 300 × g. Proteins were quantified by BCA assay (Thermo Fisher Scientific) and a total of 50 μg protein extract were used per compound treatment. Lysates were treated with vehicle (DMSO) or KBO3 (Sigma; 500 µM) for 1 hr, followed by 1 mM DBIA treatment for 1hr. Following DBIA incubation, lysates were reduced with 5 mM tris(2-carboxyethyl)phosphine hydrochloride (TCEP) (Sigma-Aldrich) for 2 min at room temperature, followed by alkylation using 20 mM chloroacetamide (Sigma-Aldrich) for 30 min in the dark at room temperature. Proteins were precipitated using SP3 magnetic beads. In brief, SP3 magnetic beads (Cytiva) were prewashed with LC-MS grade water (Sigma Aldrich), and 250 μg of combined SP3 beads (1:1, hydrophobic:hydrophilic) and LC-MS grade ethanol (Sigma-Aldrich) were added to each sample to reach a final concentration of 50% ethanol. SP3 incubation was performed for 30 min at room temperature, and beads were subsequently washed 3 times with 80% HPLC-grade ethanol (Sigma Aldrich) and then resuspended with 175 uL of Trypsin/Lys-C (1 μg, Thermo Fisher Scientific) in 200 mM EPPS (Sigma Aldrich) pH 8.4, 5 mM CaCl2. Proteins were digested overnight (16 h) at 37°C and digested peptides were enriched with streptavidin magnetic beads (Thermo Fisher Scientific) for 1 hr at room temperature. Beads were subsequently washed three times with DPBS, twice with HPLC grade-water (Sigma Aldrich). Peptides were eluted with 50% acetonitrile (Sigma Aldrich), 0.1% formic acid (Thermo Fisher Scientific), and dried using a Speedvac (Thermo Fisher Scientific). Cysteine-enriched peptides were reconstituted with 30 % acetonitrile, 70 % 200 mM EPPS pH 8.4 and labeled with 25 μg of TMT reagent (Thermo Fisher Scientific) per channel for 75 min at room temperature with rotation. Labeling was terminated by the addition of 5% hydroxylamine (Acros Organics) for 15 min followed by addition of 10% formic acid. Peptides were desalted with stage tips using the following procedure: peptides were reconstituted with 5% acetonitrile/0.1 % formic acid and loaded onto C18 Micro Spin columns(Nest Group) pre-equilibrated with LC-MS grade methanol (Fisher Chemical) and LC/MS-grade water containing 0.1% formic acid. C18 spin columns were washed 10 times with LC/MS grade water containing 0.1 % formic acid and subsequently eluted with 80% acetonitrile, 0.1% formic acid and dried using a Speedvac (Thermo Fisher Scientific).

#### Cysteine-Accessibility Proteomics Data Analysis

Peptide-level TMT reporter intensities were used to calculate, for each cysteine-containing peptide, a cysteine reactivity/accessibility value for each genotype (mean of *n* = 3 biological replicates). For each cysteine site, the change in accessibility relative to control was calculated as Δ = genotype − sgNT (percentage points; ΔsgGYS1, ΔsgPYG, ΔsgGAA), and values from peptides mapping to the same gene and cysteine were averaged. Because only triplicate-averaged values were carried forward, per-site changes are reported as point estimates and were not subjected to formal per-site significance testing.

For nucleotide-biosynthesis analyses, cysteine sites were annotated to a curated panel of 61 purine, pyrimidine, and shared-precursor genes, which were further grouped into four supply inputs: one-carbon supply (MTHFD1, MTHFD1L, MTHFD2, SHMT1, SHMT2), PRPP/ribose-phosphate supply (PRPS1, PRPSAP1, PRPSAP2), aspartate/transamination supply (GLS, GLS2, GOT1, GOT2, ADSL, ADSS1, ADSS2, PAICS, CAD), and glutamine amidotransferase supply (PPAT, PFAS, GMPS, CTPS1, CTPS2). Concordance between sgPYG and sgGAA within each category was quantified as the squared Pearson correlation (*r*^2^) between ΔsgPYG and ΔsgGAA across all sites in that category (**Figure 4E**). For the pathway diagrams, the site displayed for each enzyme was the top-changed site, defined as the cysteine with the largest |ΔsgPYG| or |ΔsgGAA| among all sites quantified for that gene (**Figure 4F** and **Figure 5C**). Sites diverging between glycogen-synthesis and glycogenolysis loss were defined by GYS1 divergence = ΔsgGYS1 − mean(ΔsgPYG, ΔsgGAA), and sites in the top decile of |GYS1 divergence| (≥11.1 percentage points) were flagged as GYS1-divergent (**Figure 5B–C**).

For visualization, cysteine sites were embedded in two dimensions using UMAP (umap-learn v0.5.12, Python) and colored by Δ accessibility for each genotype; sites belonging to the gene category of interest are circled and the largest changes labeled. Keratins (KRT1, KRT5, KRT6C, KRT9, KRT10, KRT14 and KPRP) were removed as likely contaminants, and repeated measurements of the same cysteine site were averaged. The feature matrix comprised every cysteine site quantified in all genotypes, described by its accessibility value in each genotype (galactose transition: sgNT, sgPYG and sgGAA, 25,083 sites; glucose: sgNT, sgGYS1, sgPYG and sgGAA, 21,801 sites) and z-scored per genotype. UMAP was run with n_neighbors = 15, min_dist = 0.1, Euclidean distance, two components, random initialization and random seed 42, which returns an identical embedding on repeated runs. Within each dataset a single embedding was used for all genotypes, with points colored by Δ accessibility relative to sgNT. Concordance between sgPYG and sgGAA within each supply category was calculated as the squared Pearson correlation (r²) of their Δ accessibility across the cysteine sites in that category. Structural mapping of CAD cysteines used the AlphaFold model of human CAD (UniProt P27708). Analyses were performed in Python (pandas, NumPy, SciPy, umap-learn, scikit-learn, matplotlib).

#### UMPS Immunofluorescence and Puncta Quantification

SKOV3 sgNT, sgGYS1, sgPYG, and sgGAA cells were cultured in glucose or galactose for 5 h, fixed and stained as described above with anti-UMPS (UMPS Antibody A-9, Santa Cruz Biotechnology, sc-398086) detected with an Alexa Fluor 488 secondary antibody, anti-CAD detected with an Alexa Fluor 568 secondary antibody, and DAPI. Z-stacks were acquired on a Zeiss LSM 880 confocal microscope at low exposure, using identical acquisition settings across conditions (3–4 fields per genotype and condition; 25 fields in total).

Puncta were quantified in Python from the native .czi files (aicspylibczi), reading per-file voxel dimensions from the file metadata and identifying channels by their fluorophore metadata (UMPS, Alexa Fluor 488; nuclei, DAPI). To isolate compact punctate UMPS signal from diffuse cytoplasmic staining, each Z-slice was filtered with a difference-of-Gaussians (DoG) filter (σ = 2 px minus σ = 10 px, negative values clipped to 0). A single fixed absolute DoG intensity threshold (538), calibrated once on a reference galactose image, was applied to all images so that puncta burden was directly comparable across images and conditions; objects smaller than 3 voxels were discarded. Touching puncta were separated into discrete 3D objects by a local-maxima-seeded watershed (minimum peak spacing 0.9 µm), and the equivalent spherical diameter of each object was calculated as d = (6V/π)^1/3^. Nuclei were segmented from DAPI maximum-intensity projections (Gaussian smoothing σ = 2 px, Otsu threshold, morphological opening and hole filling, distance-transform watershed with a minimum peak distance of 15 px; objects <25 µm^2^ excluded). In the absence of a membrane marker, cell territories were assigned by nucleus-seeded Voronoi tessellation of the full field. The primary readout, puncta burden, was defined as the total DoG-positive voxel volume (µm^3^) within each cell territory; discrete puncta counts and counts of large puncta (equivalent diameter ≥1.3 µm and ≥2.0 µm) were also recorded. In total, 291 cells and 1,356 puncta were analyzed (scikit-image, SciPy, NumPy, pandas).

#### Confocal Line-Scan Colocalization

For glycogen–P5CS–mitochondria colocalization (**Figure S5E**), cells were stained for glycogen (Alexa Fluor 568), mitochondria (MitoTracker Deep Red FM; Thermo Fisher Scientific, Cat# M22426; Alexa Fluor 647 channel), and P5CS (rabbit anti-ALDH18A1; Sigma-Aldrich, Cat# HPA008333; EGFP channel) and imaged by confocal microscopy. Intensity profiles along a line drawn across representative cells were extracted for each channel and normalized to the maximum intensity along the line (0–1) for display.

#### siRNA Knockdown

CAD was silenced with siRNA targeting CAD (siCAD; Horizon Discovery, L2-009471-01-0005) or a non-targeting control siRNA (siNT), transfected using Lipofectamine RNAiMAX (Thermo Fisher Scientific) as previously described^6^. Knockdown was confirmed by immunoblotting (**Figure S7C**). Cell death was measured by real-time imaging as described below, and polar metabolites were harvested as described above.

#### Real-Time Measurement of Cell Death

For temporal dynamics of cytotoxicity, cells were imaged using the CellCyte X live-cell imaging system (Discover Echo). Cells were seeded in 96-well plates, allowed to attach overnight, and transfected or treated as indicated. Cells were then incubated for 15 min with 1× CellTox Green Cytotoxicity reagent (Promega, G8731) in medium and placed into the CellCyte imaging system. Images were acquired with the 10× objective (2 images per well) every 4 h. Dead-cell fluorescence was quantified as total green-channel (CellTox Green) fluorescence in the CellCyte software using the following analysis settings: contrast sensitivity 32 a.u., separation sensitivity 50 a.u., filled hole size 50 µm^2^, and minimum object size 50 µm^2^. Data were plotted as mean ± SEM.

#### Spatial Transcriptomics Analysis

The Visium HD Spatial Gene Expression dataset (Human Lung Cancer, Breast Cancer, and Colon Cancer Tissue Array [FFPE], 11 mm Capture Area), preprocessed with Space Ranger 4.1.0, was downloaded from 10x Genomics. Data were explored and analyzed using Loupe Browser 8 (10x Genomics). A glycogenolysis gene signature (PYGL, PYGB, GAA, AGL) was generated, and differential expression analysis was performed comparing this gene set against the full dataset to identify the genes most correlated with the glycogen signature. The top 100 genes were uploaded to Enrichr to perform gene-set enrichment analysis, and the HumanCyc 2016 dataset was used to define enriched metabolic pathways.

#### Mouse Xenograft Experiment

Tumor growth experiments were performed at Beth Israel Deaconess Medical Center under the same IACUC approval described above. SUM159 cells were maintained in RPMI-1640 supplemented with 10% FBS and tested negative for mycoplasma prior to injection. On the day of injection, cells were washed twice with 1× PBS, trypsinized, and counted. A total of 5 × 10⁶ cells per mouse were resuspended in 100 µL of serum-free RPMI-1640, placed on wet ice, and mixed with Matrigel (Corning, 356230) in a 1:1 (v/v) ratio before injection into the mammary fat pad of n = 4 NSG mice. Tumors were measured with calipers twice weekly (length and width) for 28 days, after which all mice were euthanized and tumors were harvested and weighed.

#### [1-¹³C]Pyruvate Hyperpolarization

Hyperpolarized [1-¹³C]pyruvate samples were generated by either dissolution dynamic nuclear polarization (dDNP), using a SPINLab polarizer (GE Healthcare), or parahydrogen-induced polarization (PHIP), using a Polaris polarizer (NVision Quantum Technologies). dDNP sample generation followed previously published protocols ^10,11,11^ and PHIP sample generation followed standard protocols established by the vendor^12^. In brief, both approaches generated injectable solutions of [1-¹³C]pyruvate in D₂O: dDNP doses were generated in 100 mM Tris buffer with 1 mM EDTA, and PHIP doses were generated in sodium phosphate buffer (∼1:2 dibasic:monobasic) with ∼270 µM EDTA. All doses were neutralized to a pH of ∼7. Hyperpolarized samples across both techniques achieved nominal liquid-state polarizations of ∼15%–25% and concentrations of ∼75–100 mM [1-¹³C]pyruvate. sgNT tumor models were assessed with both approaches and showed comparable metabolic readouts, indicating equivalent biological performance of dDNP and PHIP doses. For each experiment, both control and experimental conditions were assessed using hyperpolarized samples generated by the same approach.

#### MRI and In Vivo ¹³C Spectroscopy

For hyperpolarized MRI, a separate cohort of NSG mice (8–10 weeks of age) was implanted subcutaneously in the right flank with 5 × 10^6^ sgNT or sgPYG SUM159 cells in a 1:1 (v/v) PBS/Matrigel mixture, and tumors were allowed to grow for ∼4–5 weeks (to no more than 1,000 mm^3^) before imaging. Animal imaging experiments were reviewed and approved by the Institutional Animal Care and Use Committee at Memorial Sloan Kettering Cancer Center (protocol 13-12-019). Imaging was performed with a 3T preclinical MRI scanner using a RAPID Tx (¹H/¹³C) volume coil and Rx (¹³C) surface coil (Bruker). Prior to imaging, tumor-bearing animals were tail-vein catheterized, anesthetized with 2%–3% isoflurane, positioned on top of the surface coil with tumors centralized to the detection volume, and secured with an approved spine immobilizer to allow the animal to wake during the hyperpolarized scan. Once positioned at isocenter, anatomical images were acquired with a ¹H T₂-weighted rapid acquisition with relaxation enhancement (RARE) sequence with the following parameters: 33 ms TE, 8.25 ms echo spacing, 2,500 ms TR, 128 × 128 matrix, 40 × 40 mm FOV, 2 mm slices, 2 averages, 20 kHz bandwidth. Isoflurane administration was then discontinued, animals were allowed to wake (>120 bpm), and 200 µL of hyperpolarized [1-¹³C]pyruvate was infused over 30 seconds with a power injector. The first spectroscopy scan was acquired 15 seconds after the start of the infusion. Dynamic ¹³C spectroscopy scans were acquired as previously described^10^ with the following parameters: 60° flip angle, 2,048 points, 3-second temporal resolution, and 3,236.2 Hz spectral width. After ¹³C scans, anesthesia was resumed and confirmatory ¹H anatomical images were acquired to ensure tumor positioning had not changed. At the completion of the imaging session, animals were removed from the scanner and humanely euthanized.

All spectra were processed using MestReNova (Mestrelab Research S.L.): spectra were line-broadened, phased, baseline-corrected, zero-filled, and integrated to quantify peak areas. Peak areas were then used to calculate lactate-to-bicarbonate ratios and total-carbon-normalized values for all metabolic products.

#### MALDI Mass Spectrometry Imaging

Breast tumors from SUM159 xenograft experiment were surgically removed and slow-frozen in liquid nitrogen. Cryosections (10 µm) were cut on a Leica CM1860 cryostat at −15 °C (chilled to −20 °C before cutting), thaw-mounted without optimal-cutting-temperature compound onto Bruker IntelliSlides, and dehydrated in a vacuum desiccator for 1 h to halt metabolic activity. Four sections per genotype were distributed across four slides and two acquisition runs, with each slide carrying one NT, one PYG, and one GAA section. N-(1-naphthyl)ethylenediamine dihydrochloride (NEDC) matrix (7 mg/mL in 70% methanol) was applied with an HTX M5 sprayer (14 passes, 0.06 mL/min, 3 mm offset, 1,200 mm/min, nozzle 30 °C, 10 psi, tray 50 °C). Imaging was performed on a Bruker timsTOF fleX with acquisition set up through the flexImaging autopilot using a masked .mis template of the tissue regions (laser 10,000 Hz, 60% energy, 300 shots per pixel, 46 µm laser raster producing 50 µm pixels) in negative-ion mode across m/z 50–2,000, as described^13–16^. Features were extracted in MetaboScape (Bruker) with the T-ReX 2D algorithm (intensity threshold 500; [M−H]⁻, [M+Cl]⁻ and [M−H₂O]⁻ ions) and annotated by accurate mass within 5–10 ppm and mSigma 25–250 against HMDB, MoNA, MassBank, FiehnLib LipidBlast, MetaboBASE and NIST 2022 libraries.

#### Glycogen MALDI-MSI

Glycogen was imaged on the same sections after metabolite acquisition, using the sequential multi-omic MALDI workflow described previously ^13,15,16^. After NEDC imaging, slides were removed from the instrument, the NEDC matrix was stripped by a 5 min wash in ice-cold 100% methanol, and sections were fixed for 1 h in 10% (v/v) neutral-buffered formalin (37% formaldehyde in 1× PBS). Sections were then dehydrated for 2 h in 70% ethanol, followed by progressive 5 min washes in 70%, 80%, 90% and 100% ethanol, and desiccated overnight. Before antigen retrieval, sections were rehydrated by a 1 min wash in 70% ethanol followed by two 3 min washes in water. Antigen retrieval was performed in 0.05% (v/v) citraconic anhydride (pH 3.0 ± 0.5) at 95 °C for 30 min, after which slides were vacuum-desiccated. Isoamylase and PNGase F (3 U of each enzyme reconstituted in 1 mL HPLC-grade water) were applied with an HTX M5 sprayer (15 passes, 0.025 mL/min, 3 mm offset, 900 mm/min, nozzle 45 °C, tray at ambient temperature); isoamylase hydrolyzes the α-1,6 branch points of glycogen to release its linear α-1,4-glucan chains, and PNGase F releases N-linked glycans^17^. Slides were incubated at 37 °C for 2 h in a humidified chamber and vacuum-desiccated before matrix application. α-Cyano-4-hydroxycinnamic acid (CHCA) matrix (7 mg/mL in 50% HPLC-grade acetonitrile containing 0.01% trifluoroacetic acid) was then applied with the HTX M5 sprayer (10 passes, 0.1 mL/min, 3 mm offset, 1,200 mm/min, nozzle 79 °C, tray 50 °C). Glycogen was imaged on the timsTOF fleX in positive-ion mode across m/z 600–4,000 (laser 10,000 Hz, 70% energy, 396 shots per pixel), using the same 46 µm laser raster, 50 µm pixels and masked .mis template as the NEDC acquisition^13^. Isoamylase-released glucan chains were detected as a series of sodiated glucose polymers ([M+Na]⁺) spaced by one hexose unit (162.05 Da), for example chain length 7 at m/z 1,175.37, and were annotated by accurate mass (within 20 ppm of the theoretical [M+Na]⁺ m/z)^13,18^. Total glycogen was calculated as the summed TIC-normalized peak area of chain lengths 3–18 (m/z 527.16–2,957.95)^18^, and this summed signal is the glycogen image shown in Figures 5H and S8.

#### MALDI-MSI Data Processing and Statistics

Images were processed in SCiLS Lab (Bruker Daltonics), and pixel-level intensities for annotated metabolites were exported through the SCiLS Lab API with total-ion-current (TIC) normalization. Analyses were performed in Python 3.11 (pandas 3.0, NumPy 2.4, SciPy 1.17, statsmodels 0.15, matplotlib 3.10). Intensities were log1p-transformed (ln[1 + intensity]); zero-intensity pixels were retained. NT (n = 4 tissue sections; 57,893 pixels) and PYG (n = 4; 46,677 pixels) were compared for each metabolite with a linear mixed-effects model fitted by restricted maximum likelihood on all pixels, with genotype as a fixed effect and tissue section as a random intercept to account for spatial pseudoreplication. Significance was assessed by the Wald test on the genotype coefficient. Pyruvate, lactate, citrate, cis-aconitate, malate and glutamate were each selected a priori as individual hypotheses, and p-values are reported without correction for multiple comparisons. Fold differences were calculated as exp(Δ) of the mean per-section log1p intensity (PYG/NT). Violin plots show the distribution of all pixels, with box plots indicating median and interquartile range (whiskers, 1.5× IQR) and points indicating per-section means; n represents tissue sections.

#### MALDI-MSI Spatial Correlation Analysis

For glycogen, UMP, and aspartate co-distribution (**Figures 5H and S8**), TIC-normalized pixel intensities were exported for each annotated feature from all sgNT (sections 1–4), sgPYG (sections 9–12), and sgGAA (sections 13–16) sections. The NEDC (negative-ion) and CHCA (positive-ion) acquisitions were collected from the same sections using a single masked flexImaging .mis template, so that both datasets share a common 50-µm pixel grid^13,15^. Pixel tables for each acquisition were exported from SCiLS Lab with their x–y coordinates and co-registered section by section: coordinates were first aligned by the offset between the tissue bounding boxes in the two datasets, and pixels were then matched on the exact 50-µm grid by nearest-neighbor lookup (SciPy cKDTree), retaining only pixels present in both acquisitions (136,169 matched pixels across the 12 sections). Glycogen (CHCA) was thereby paired pixel-for-pixel with UMP and aspartate (NEDC), and all images, correlations and line scans below were derived from these matched pixels. For each section, Spearman rank correlations between each metabolite pair were calculated across all matched 50-µm pixels, and per-genotype values are reported as mean ± SD of per-section coefficients (*n* = 4 sections/genotype). Single-metabolite images were displayed with sequential color maps scaled to the indicated intensity ranges and overlaid as a three-color composite. Line-scan profiles were extracted along a transect through each section and min–max normalized for display.

### QUANTIFICATION AND STATISTICAL ANALYSIS

All data was analyzed using Graphpad Prism 11 unless otherwise stated. All quantitative data are presented as the mean ± standard deviation (SD) or mean ± standard error of the mean (SEM), as stated in each figure legend: SD was used for the in vivo xenograft (Figure 1D, E) and hyperpolarized-¹³C-MRI (Figure 1H, I) datasets, and SEM was used for all in vitro growth, metabolite-tracing, and viability datasets (**Figures 2 and 3 and their associated panels**).

For the in vivo xenograft study (**Figure 1D, E**), tumor volume and endpoint tumor weight were compared between sgNT and sgPYG tumors (n = 4 mice/group) by two-tailed unpaired Student’s t-test. In the independent hyperpolarized [1-¹³C]pyruvate MRI cohort (**Figure 1G–I; n = 5 sgNT, n = 4 sgPYG mice**), normalized lactate, alanine, and bicarbonate signal intensities, and the derived lactate-to-bicarbonate ratio, were likewise compared between genotypes by two-tailed unpaired t-test (exact P values: P = 0.02 for bicarbonate signal, P = 0.009 for the lactate-to-bicarbonate ratio).

For all in vitro comparisons across three or more groups (sgNT, sgPYG, and sgGAA, and, in **Figure 5**, sgGYS1) — including SRB growth/viability assays (**Figure 2C, G; Figure 3B, C, H**), isotope-tracing quantification (**Figure 2E, H; Figure 3A, F**), and redox-cofactor ratio measurements (**Figure 3D**) — group means were compared by one-way ANOVA with post hoc multiple-comparisons test. Where a panel displays more than one pairwise comparison, symbols distinguish which comparison is being made rather than degree of significance: in Figure 2C, asterisks (*) denote sgNT vs. sgPYG and number signs (#) denote sgNT vs. sgGAA; in Figure 2H, asterisks (*), number signs (#), and dollar signs ($) denote comparisons at the m+0, m+1, and m+2 isotopologues, respectively. An *a priori* significance threshold of α = 0.05 was applied to all statistical comparisons reported in this study; comparisons with p ≥ 0.05 are reported as not significant (ns). Across all figures, significance is denoted as *P < 0.05, **P < 0.01, ***P < 0.001, ****P < 0.0001, and ns (not significant).

Unless otherwise noted, n reflects biological replicates: independently generated CRISPR-edited cell-line pools for cell-based assays, independent tumors/animals for in vivo studies, or independently harvested plates/dishes for metabolomics and proteomics (n = 3 biological replicates for all polar metabolomics, isotope-tracing, and cysteine-accessibility proteomics experiments, per Method Details).

For UMPS puncta quantification (**Figure 5I**), puncta burden is plotted per cell (violin plots); because cell number per field ranged from 2 to 30, condition-level summaries were also calculated as the mean and median of per-field means to avoid overweighting fields with many segmented cells. Per-cell puncta burden was compared across genotypes and carbon sources by two-way ANOVA with Tukey’s multiple comparisons test. All fields were retained for analysis, including one sgNT galactose field in which UMPS signal was more diffuse than punctate.

For MALDI-MSI spatial correlation (**Figures 5H and S8**), Spearman correlation coefficients were calculated per section across all matched pixels and summarized as mean ± SD across sections (*n* = 4 sections/genotype); no formal between-genotype test was applied.

For the cysteine-accessibility (DBIA/TMT) and whole-cell proteomics datasets (**Figures 3E, 4D– F, 5B–D**), per-site or per-protein accessibility/abundance changes are reported as fold-change or Δaccessibility relative to sgNT. Concordance between the sgPYG and sgGAA accessibility signatures across the four nucleotide-biosynthesis input categories (**Figure 4E**) is reported as r² per category using Pearson correlation coefficient. The volcano plot of CBM20 pulldown metabolites (**Figure 5G**) reports log₂ fold-change vs. −log₁₀ P value, calculated using two-tailed t-test across 3 biological replicates.

