## Supplementary figures for "Glycogen fuels and spatially regulates nucleotide biosynthesis in cancer"

**Figure S1. Loss of cytoplasmic glycogenolysis restrains tumor growth and TCA cycle metabolite abundance in vivo, related to Figure 1**

**(A)** Immunoblot of PYGB in sgNT and sgPYG SUM159 cells used for xenografts. Vinculin, loading control.

**(B)** Xenograft design. sgNT or sgPYG SUM159 cells ($5 \times 10^6$ per mouse) were implanted into the mammary fat pad of NSG mice (n = 4 per group), measured by caliper on days 7, 14, 22, 25, and 28, and harvested on day 30.

**(C and D)** Tumor volume over time (C) and tumor weight at harvest (D).

**(E)** Representative H&E staining of tumors from three mice per group; insets, low magnification. Scale bar, 50 μm.

**(F)** Total-carbon-normalized lactate, alanine, and bicarbonate signal from hyperpolarized [1-$^{13}$C]pyruvate spectroscopy of subcutaneous xenografts (n = 5 sgNT and 4 sgPYG mice).

**(G)** MALDI-MSI of the metabolites in Figure 1E across all four sgNT and sgPYG sections. Scale bar, 7 mm.

Data are represented as mean ± SD; two-tailed unpaired t test (*$p < 0.05$, **$p < 0.01$; ns, not significant).

**Figure S2. Cytoplasmic and lysosomal glycogenolysis are required for growth under galactose stress across cancer cell lines, related to Figure 2**

**(A)** Immunoblots of PYGB, PYGL, and GAA in SKOV3 and SUM159 cells. Vinculin, loading control.

**(B)** Growth of PC3 (top) and 143B (bottom) cells in 10 mM glucose (left) or 10 mM galactose (right), measured by SRB absorbance. * and # denote sgNT versus sgPYG and sgNT versus sgGAA, respectively.

**(C)** Immunoblots of PYGB, PYGL, and GAA in PC3 and 143B cells. Vinculin, loading control.

**(D)** Relative living cell number of SKOV3 cells in glucose (+Gluc), glucose-free (−Gluc), or galactose (+Gal) medium.

**(E)** Relative living cell number of SKOV3 cells in response to the glycogen phosphorylase inhibitor CP91149 in 10 mM glucose or 10 mM galactose.

**(F)** Relative living cell number of SKOV3, SUM159, PC3, A549, 143B, and HeLa cells treated with DMSO, the GLUT1 inhibitor BAY-876 (BAY; 1 μM), CP91149 (CP; 50 μM), or both.

Data are represented as mean ± SEM; n = 3 biological replicates (n = 2 in F); one-way ANOVA with multiple comparisons (**p* < 0.05, ***p* < 0.01, ****p* < 0.001, *****p* < 0.0001; ns, not significant).

**Figure S3. Glycogen is mobilized within 1 h of galactose stress, and glycogen phosphorylase inhibition phenocopies genetic loss of glycogenolysis in TCA cycle labeling, related to Figure 2**

**(A)** Top altered polar metabolites in SUM159 cells after 1 h in galactose (row normalized, hierarchical clustering).

**(B)** Representative immunofluorescence of nuclei (DAPI, blue), F-actin (phalloidin, green), mitochondria (MitoTracker, red), and glycogen (magenta) in SKOV3 (top) and SUM159 (bottom) cells in glucose or after 1 or 6 h in galactose. Scale bar, 20 μm.

**(C)** Glycogen pulse-chase in SUM159 cells, as in Figure 2E.

**(D–F)** Glycogen phosphorylase inhibitor chase. SKOV3 (D), SUM159 (E), and A549 (F) cells were labeled with U-$^{13}$C-glucose for 24 h and then treated with DMSO or CP91149 in unlabeled medium for 6 h (top). Label incorporation into PEP m+3 and citrate, glutamate, malate, OAA, and aspartate m+2 is shown below.

Data are represented as mean ± SEM; n = 3 biological replicates; one-way ANOVA (C) or two-tailed unpaired t test (D–F) (**p* < 0.05, ***p* < 0.01, ****p* < 0.001, *****p* < 0.0001; ns, not significant).

**Figure S4. Glycogenolysis loss alters pyruvate carbon fate and the malate/citrate labeling ratio and blunts rescue by NAD(P)H reoxidation, related to Figures 2 and 3**

**(A)** Top altered $^{13}C$ isotopologues in cells cultured in galactose with U-$^{13}C$-pyruvate, as in Figure 2H (row normalized, hierarchical clustering).

**(B)** m+2 malate/m+2 citrate labeling ratio in SUM159 cells, from the tracing in Figure S3C.

**(C)** m+2 malate/m+2 citrate labeling ratio in SKOV3, SUM159, and A549 cells treated with DMSO or CP91149, from the tracing in Figures S3D–S3F.

**(D)** Relative viability in galactose of cells expressing the NADH- or NADPH-reoxidizing enzymes AOX, LbNOX, or mitoLbNOX.

(E) Pentose phosphate pathway intermediates in SKOV3 sgNT, sgPYG, and sgGAA cells labeled overnight with U-$^{13}C$-glucose and chased into unlabeled galactose for 6 h, showing isotopologue labeling and total pool size.

Data are represented as mean ± SEM; n = 3 biological replicates; one-way ANOVA (B, D, and E) or two-tailed unpaired t test (C) (*$p < 0.05$; ns, not significant).

**Figure S5. Glycogenolysis loss raises the NADP$^+$/NADPH ratio and impairs reductive glutamine metabolism in vitro and in vivo, related to Figure 3**

**(A and B)** NAD$^+$/NADH (left) and NADP$^+$/NADPH (right) peak-area ratios from polar metabolomics of SKOV3 (A) and SUM159 (B) cells after 6 h in galactose.

**(C)** U-$^{13}C$-glutamine tracing (8 h) in SUM159 cells in galactose, as in Figure 3F.

**(D)** MALDI-MSI of proline (*m/z* 114.0559), ornithine (*m/z* 131.0821), and malic acid (*m/z* 133.0137) in all sgNT, sgPYG, and sgGAA SUM159 xenograft sections. Scale bar, 7 mm.

**(E)** Representative confocal images of SKOV3 cells stained with MitoTracker (gray), glycogen (red), and P5CS (green) (left; scale bar, 10 μm), with line-scan intensity profiles from representative cells (right).

Data are represented as mean ± SEM; n = 3 biological replicates; one-way ANOVA with multiple comparisons (*$p < 0.05$, **$p < 0.01$, ***$p < 0.001$, ****$p < 0.0001$; ns, not significant).

**Figure S6. Glycogenolysis loss perturbs nucleotide pools and labeling and differentially alters sensitivity to CTPS1 and GOT1 inhibition, related to Figure 4**

**(A)** Top altered polar metabolites in SKOV3 cells after 6 h in galactose (row normalized, hierarchical clustering).

**(B)** Amide-$^{15}$N-glutamine-labeled isotopologues of nucleotides and related metabolites in SKOV3 cells labeled for 12 h in galactose.

**(C)** U-$^{13}$C-glucose-labeled isotopologues of nucleotides and related metabolites in SKOV3 cells labeled for 24 h and chased into unlabeled galactose for 6 h.

(D) UTP, glyceraldehyde-3-phosphate, dihydroxyacetone phosphate, and sn-glycerol-3-phosphate in sgNT, sgPYG, and sgGAA cells treated with DMSO or dencatistat for 24 h.

**(E)** Aspartate, α-ketoglutarate, glutamate, and oxaloacetate in cells treated with DMSO or 10 μM GOT1-IN-1 for 24 h (schematic, left).

Heatmaps in (A)–(C) are row normalized with hierarchical clustering. Data in (D) and (E) are represented as mean ± SEM; n = 3 biological replicates; one-way ANOVA with multiple comparisons (*$p < 0.05$).

**Figure S7. Glycogen synthesis loss creates a distinct pyrimidine dependency, and CBM20 glycogen immunoprecipitation captures glycogen-associated proteins and pyrimidine intermediates, related to Figure 5**

**(A)** Immunoblots of GYS1, PYGB, PYGL, and GAA in SKOV3 sgNT, sgGYS1, sgPYG, and sgGAA cells. Actin, loading control.

**(B)** Relative growth in galactose with or without 100 μM uridine.

**(C)** Immunoblot of CAD in siNT- and siCAD-transfected cells. Vinculin, loading control.

**(D)** UMP, UTP, CMP, and CTP in siNT- and siCAD-transfected cells.

**(E)** Immunoblots of input (pre-IP), flowthrough, and anti-FLAG immunoprecipitates from HEK293-CBM20-FLAG cells lysed in KPBS or NP-40 buffer and immunoprecipitated for 15 min or 4 h, probed for PYGB (~97 kDa), glycogen synthase (GS; 84 kDa), GAA (105 kDa), and actin (45 kDa).

**(F)** Top differentially bound metabolites in CBM20 immunoprecipitates from sgNT versus sgGYS1 cells (row normalized, hierarchical clustering).

**(G)** Orotidine-5′-phosphate, PRPP, UTP, and aspartate in CBM20 immunoprecipitates from sgNT, sgPYG, and sgGAA cells.

Data are represented as mean ± SEM; n = 3 biological replicates; one-way ANOVA with multiple comparisons (*$p < 0.05$, **$p < 0.01$, ***$p < 0.001$, ****$p < 0.0001$; ns, not significant).

**Figure S8. Spatial co-distribution of glycogen with UMP and aspartate in xenograft tumors, related to Figure 5**

**(A)** MALDI-MSI of glycogen, UMP, and aspartate, and their composite, in all sgNT (sections 1–4), sgPYG (sections 9–12), and sgGAA (sections 13–16) SUM159 xenograft sections. Per-section pixel Spearman correlations are shown beside each row.

**(B)** Line-scan profiles of glycogen, UMP, and aspartate intensity in each section, with the per-section pixel Spearman *r* (as in C).

**(C)** Pixel Spearman correlations per section across all matched 50-μm pixels, with mean ± SD per genotype (n = 4 sections per genotype).

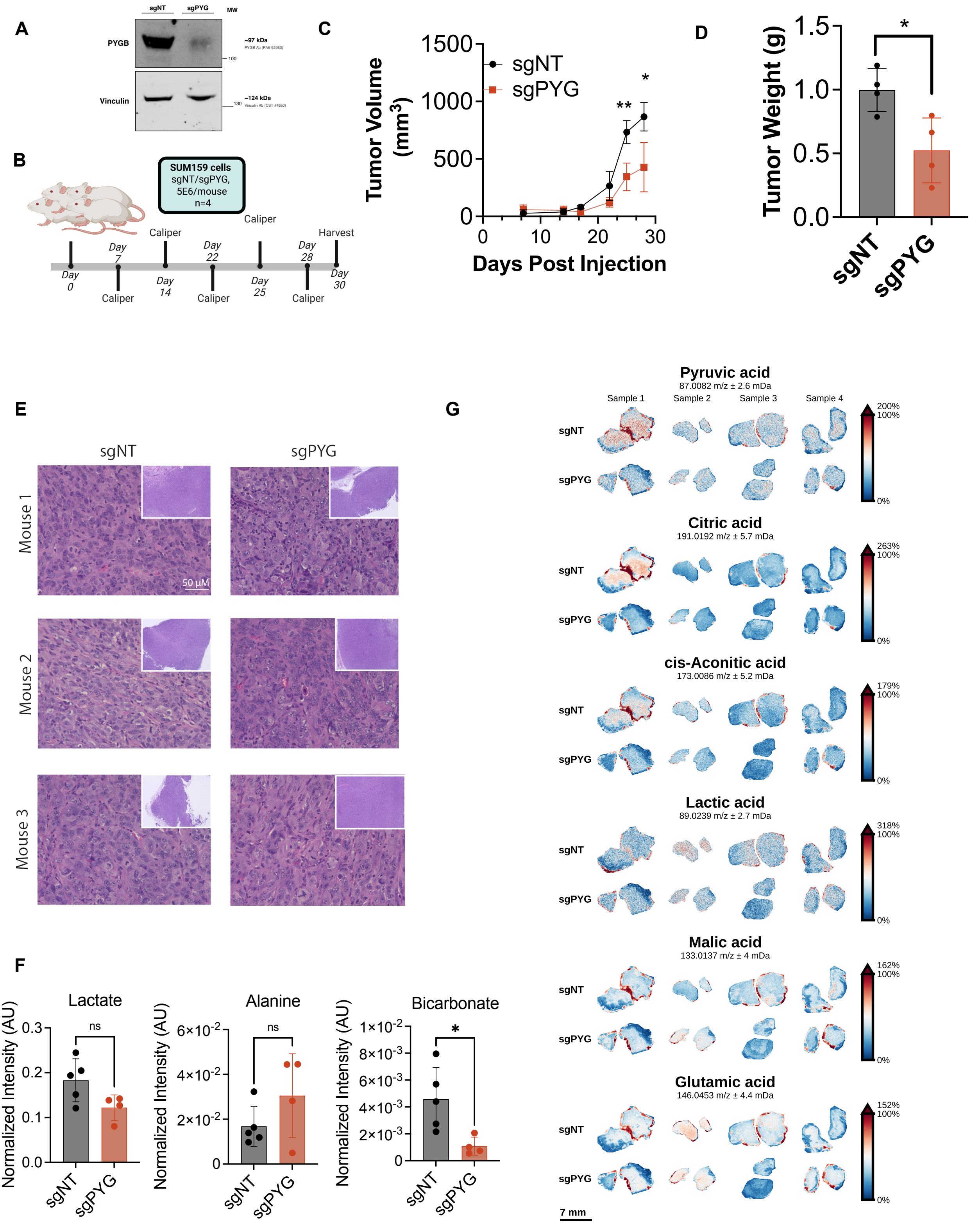


Khan et al, Figure S1

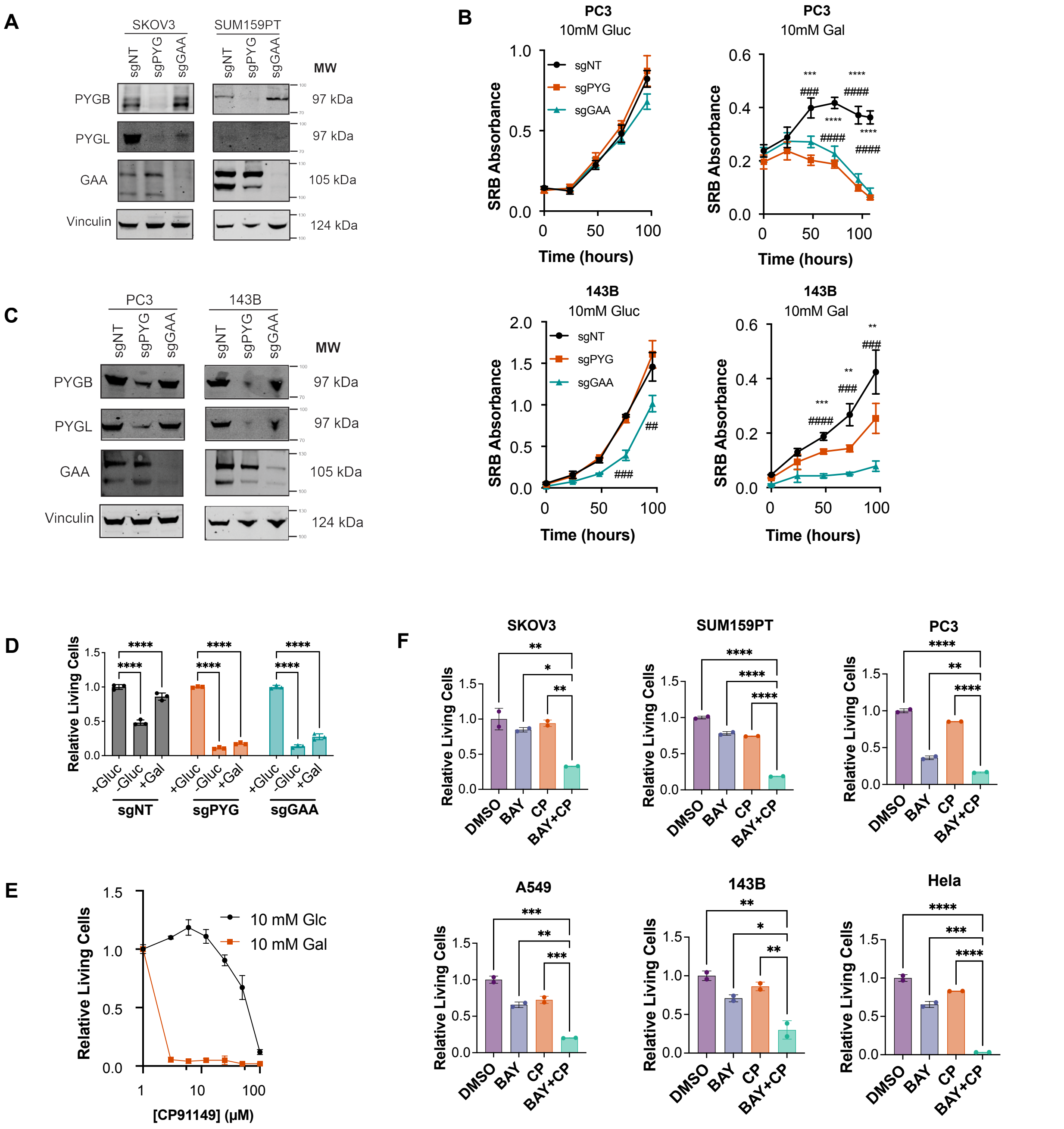


Khan et al, Figure S2

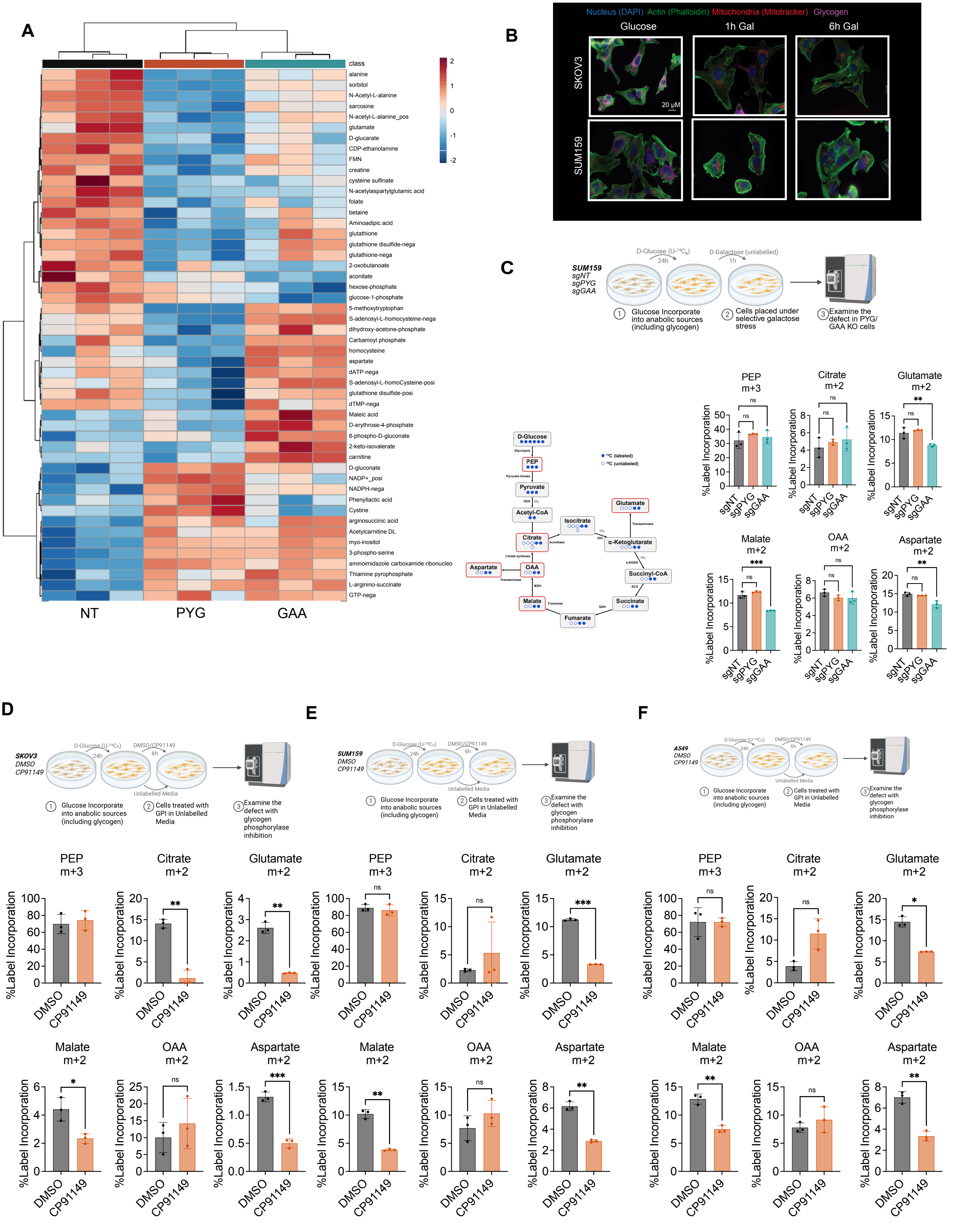


Khan et al, Figure S3

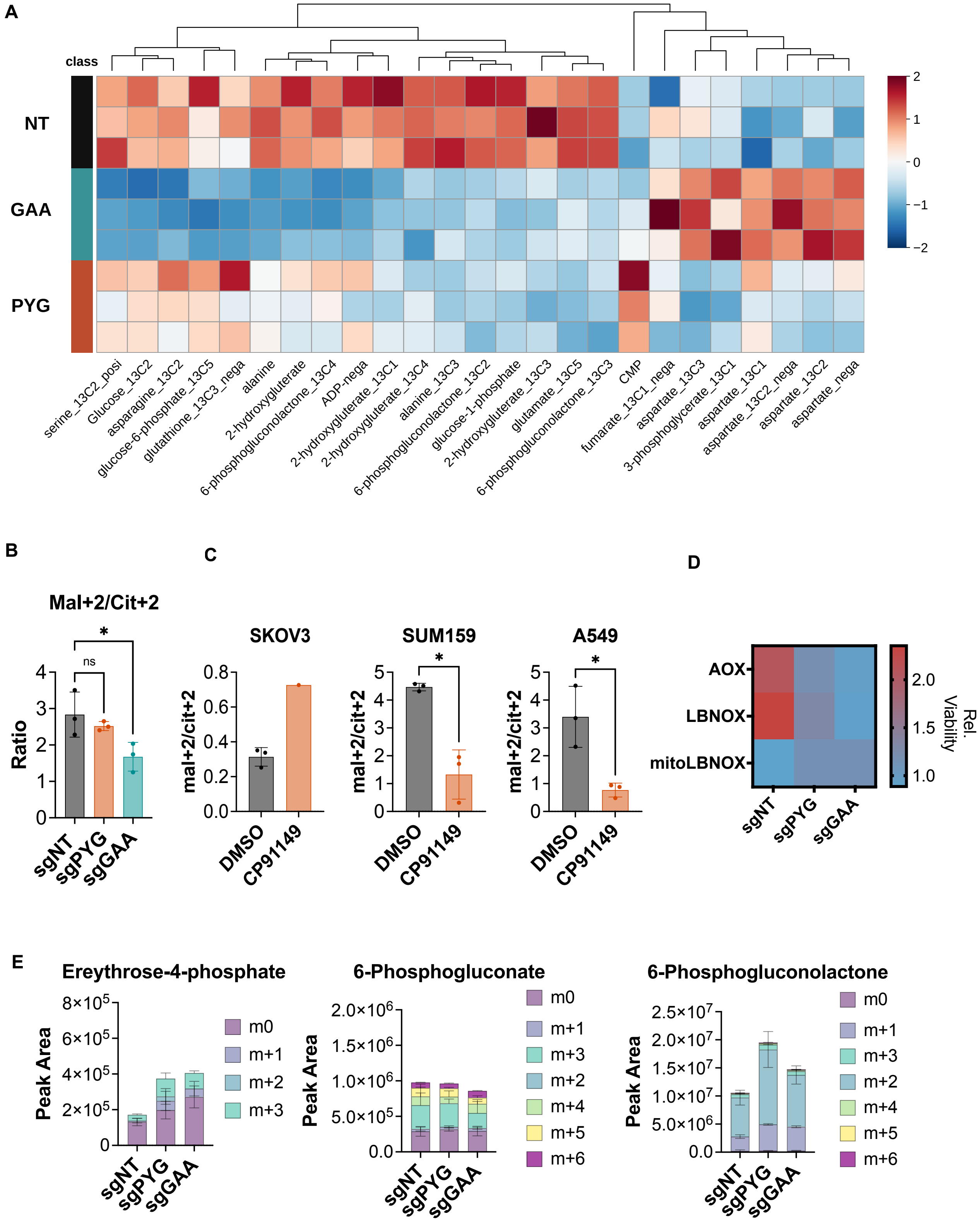


Khan et al, Figure S4

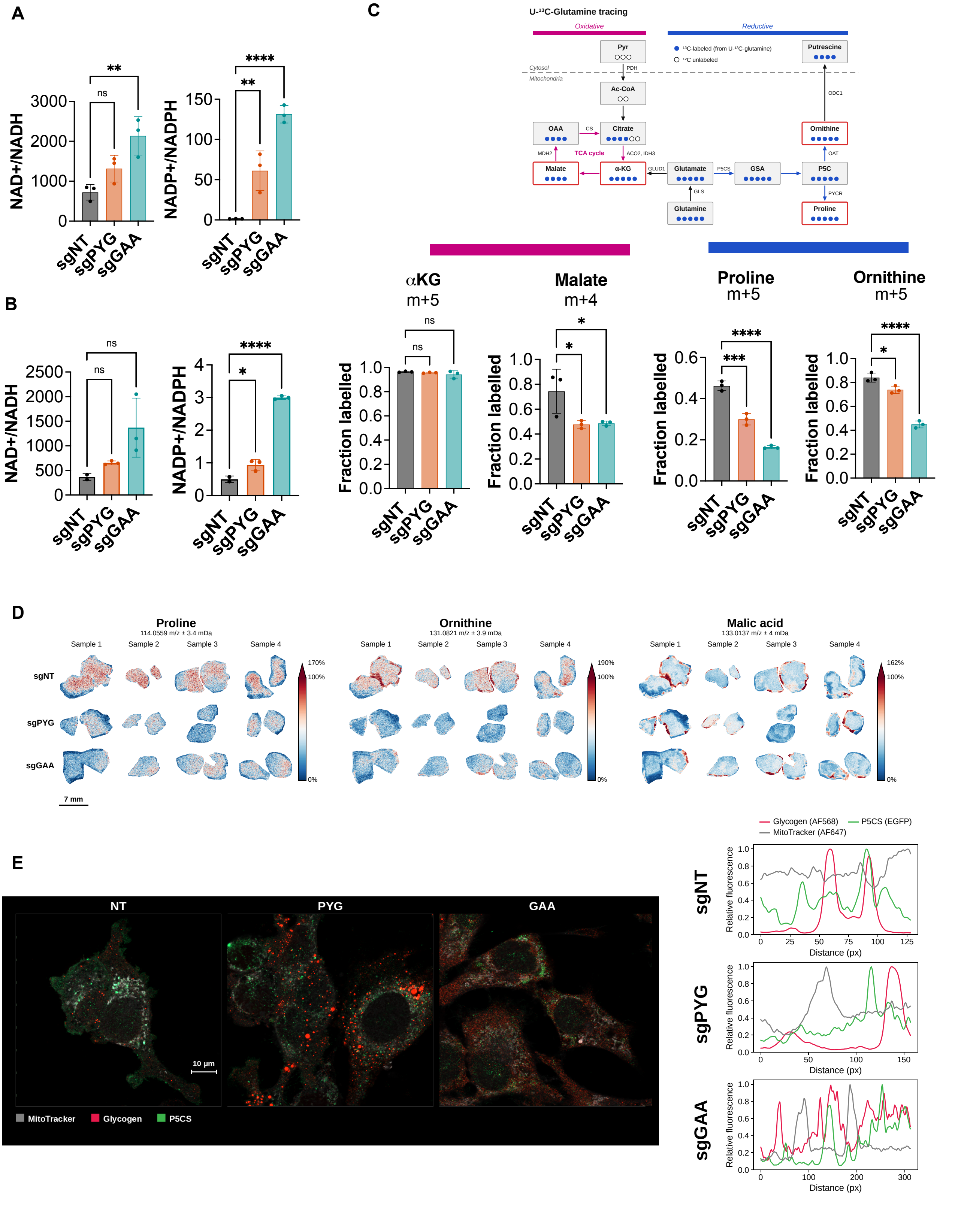


Khan et al, Figure S5

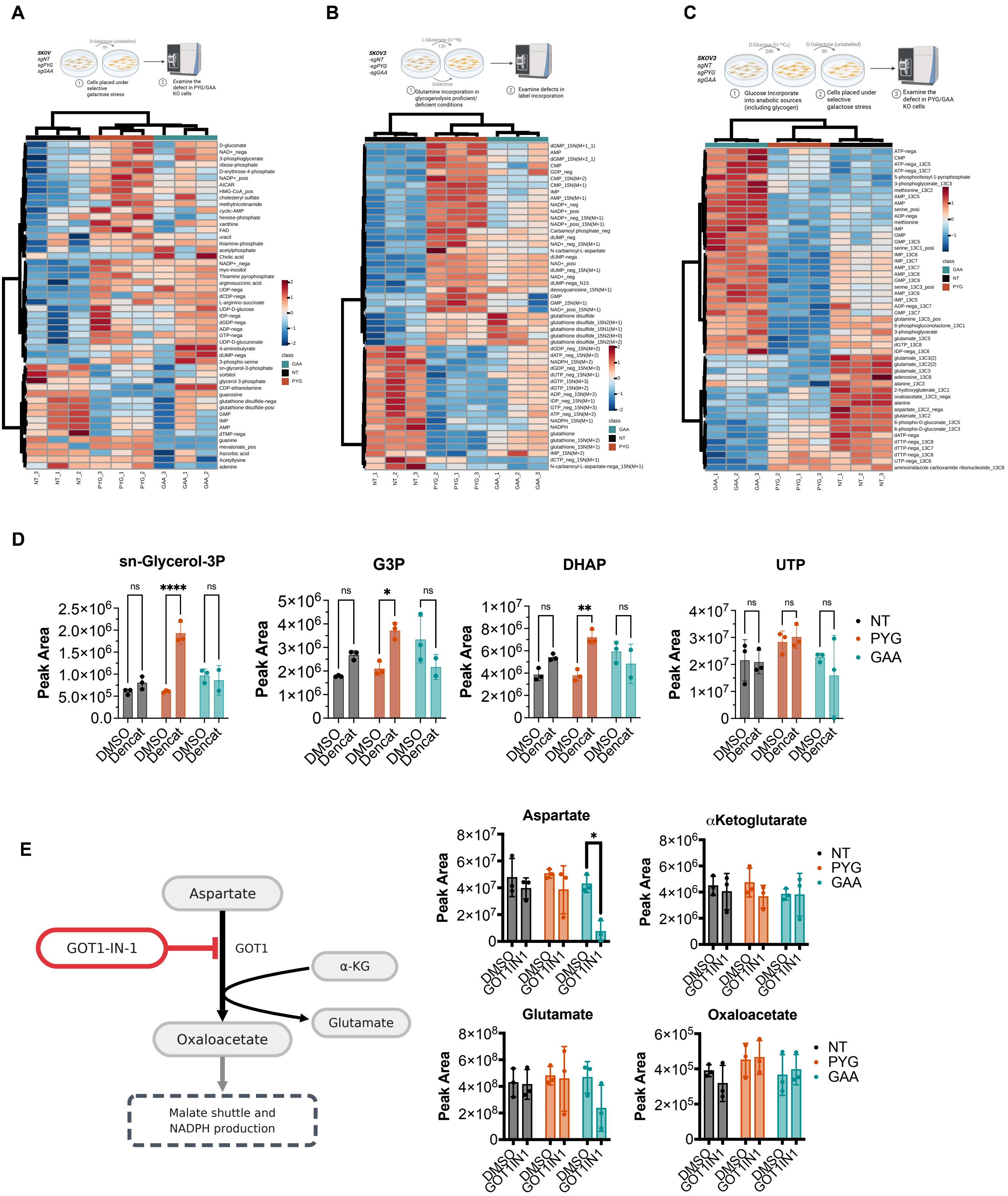


Khan et al, Figure S6

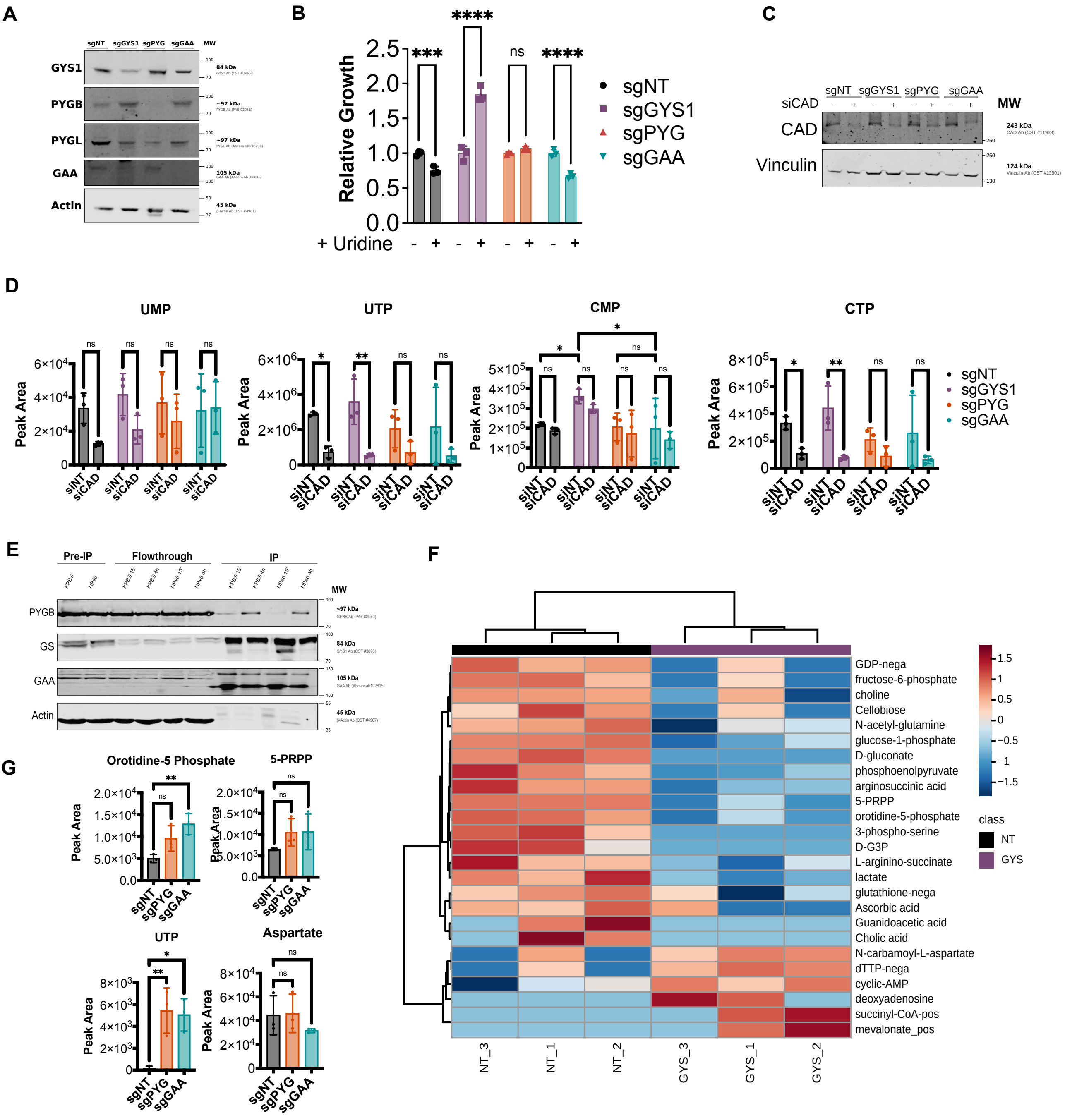


Khan et al, Figure S7

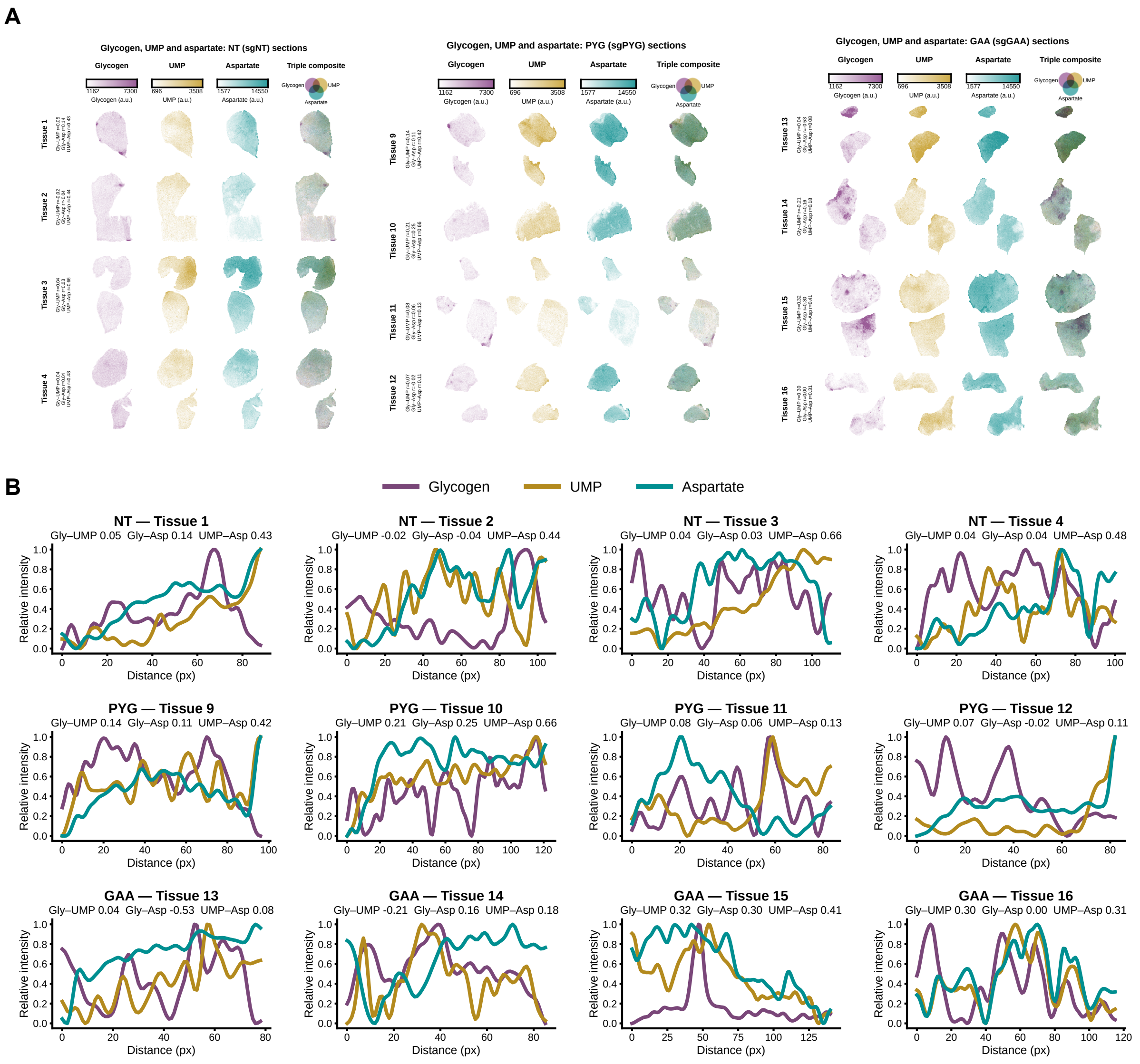


**C**

### Pixel correlation per section (Spearman r)

| | sgNT | | | | sgPYG | | | | sgGAA | | | | Mean ± SD | | |
|---|---|---|---|---|---|---|---|---|---|---|---|---|---|---|---|
| **Tissue** | 1 | 2 | 3 | 4 | 9 | 10 | 11 | 12 | 13 | 14 | 15 | 16 | sgNT | sgPYG | sgGAA |
| **Glycogen–UMP** | 0.05 | −0.02 | 0.04 | 0.04 | 0.14 | 0.21 | 0.08 | 0.07 | 0.04 | −0.21 | 0.32 | 0.30 | **0.03 ± 0.03** | **0.12 ± 0.06** | **0.11 ± 0.25** |
| **Glycogen–Aspartate** | 0.14 | −0.04 | 0.03 | 0.04 | 0.11 | 0.25 | 0.06 | −0.02 | −0.53 | 0.16 | 0.30 | 0.00 | **0.04 ± 0.07** | **0.10 ± 0.12** | **−0.02 ± 0.36** |
| **UMP–Aspartate** | 0.43 | 0.44 | 0.66 | 0.48 | 0.41 | 0.66 | 0.13 | 0.12 | 0.08 | 0.18 | 0.41 | 0.30 | **0.50 ± 0.10** | **0.33 ± 0.26** | **0.24 ± 0.14** |

*Spearman r across all matched pixels in each section (50 μm). Mean ± SD, n = 4 sections per genotype.*
